# Mathematical model of phenotypic plasticity predicts interconversion is a drug target for neuroblastoma

**DOI:** 10.64898/2026.11.04.764313

**Authors:** Francesca Covell, Matishalin Patel, Kenneth.Y. Wertheim

## Abstract

Neuroblastoma is a paediatric cancer arising from the sympathetic nervous system as neural crest cells, made up of noradrenergic (NOR/ADRN/N), mesenchymal like (MES/S), and intermediate (I-type/I) cell types. These cells can interconvert between types, contributing to treatment resistance and relapse. We developed a mathematical model to investigate how interconversion patterns affect the efficacy of phenotype-specific inhibitors. We used three ordinary differential equations to describe the dynamics of the three populations, including logistic growth, interconversion, and drug-induced death. Of the 64 possible interconversion patterns, five reduced the target population: two for N cells and three for S cells. N targeting patterns were more effective with higher interconversion rates, while mesenchymal targeting patterns were more effective with lower interconversion rates. Stability analysis revealed that natural tumour growth was always unstable, but during treatment, N-suppressing patterns combined with an S inhibitor gave the most consistently stable origin. Testing multi-stage and cyclic treatment schedules revealed that N-suppressing patterns with an S inhibitor, followed by an S-suppressing pattern with an N inhibitor, achieved more reliable remission. Three of the five patterns of interest correspond to reported biological plausibility, and known compounds exist to induce most of the required transitions, supporting the clinical relevance of these strategies for improving neuroblastoma treatment outcomes.

## 1 Introduction

Neuroblastoma is a paediatric cancer arising from the sympathetic nervous system as neural crest cells. These cells differentiate during which genomic or epigenetic changes occur leading to neuroblastoma [1, 2]. High-risk neuroblastoma patients have a longterm survival rate below 50% even after multi-modal therapy. Solid neuroblastoma tumours are composed of three major cell types [2–4].

Standard treatment begins with induction chemotherapy and surgery; followed by a consolidation phase involving high-dose chemotherapy and radiation therapy; and a maintenance phase involving anti-GD2 immunotherapy [5, 6]. Beyond the standard protocol, targeted therapies for neuroblastoma have emerged and are being actively researched [7]. The three cell types have been shown to be resistant to different treatments [8–10].

Different neuroblastoma cell lines contain varying proportions of the three cell populations [4, 11, 12]. Two of these cell types— noradrenergic (NOR) and mesenchymal (MES)—are genetically clonal but epigenetically distinct [4, 8, 13, 14]. The third, an intermediate (I-type) population, has been described as detrimental to treatment outcomes in neuroblastoma [3, 15].

The terms noradrenergic (NOR) and adrenergic (ADRN) have been used interchangeably in the literature [8]. Because the cells in neuroblastoma produce noradrenaline, we use NOR rather than ADRN throughout this paper [4, 8, 14]. In addition, the existence of I-type cells within cell lines is still under investigation [14].

Noradrenergic cells are the most rapidly proliferating of the three types [16, 17] and express neuronal differentiation markers, including sympathoadrenergic lineage genes and neurotransmitter-associated enzymes [18]. Despite their proliferative nature, neuroblastoma dominated by noradrenergic cells have led to more favourable clinical outcomes. This is likely due to noradrenergic phenotypes resembling normal neuroblasts, which correlates with a positive treatment response, where as relapsed, chemoresistant tumours show higher immature mesenchymal-like populations[19]. Mesenchymal cells are less differentiated than noradrenergic cells, resemble neural crest progenitor cells and are more migratory leading to a higher chance of metastasis [8, 13, 18]. Papers that suport the existance of I-type cells discribe an intermediate morphology that contribute to tumorigenesis and heterogeneity, associated with poor clinical outcomes [3, 15, 20, 21].

Different neuroblastoma patient-derived cell lines, xenografts, and patient samples contain varying proportions of the three cell populations [4, 11, 12, 22]. Populations at diagnosis are: noradrenergic 40–60%, mesenchymal 10% [23]. This tumour heterogeneity is associated with treatment failure and relapse [2, 24, 25]. Additionally,neuroblastoma cells can transdifferentiate between the three types through core regulatory circuitries (CRCs) composed of transcription factors [2, 8, 13, 26, 27], allowing escape from cell type–specific inhibitors.

There are three cell types within the neuroblastoma tumour, with six possible direct pathways linking pairs of these types. Several mechanisms have been proposed to explain how the three cell types interconvert, including stochastic interconversion between the three types [2, 13, 18, 27], bidirectional noradrenergic and mesenchymal via I-type [18], and clonal competition (no direct interconversion) [28]. If each of the six pathways is treated as binary (active or inactive), there are 2^6^ = 64 possible combinations, which will henceforth be described as interconversion patterns. Of these 64 patterns 34 are deemed plausible for natural occurrence [18].

Because neuroblastoma tumours comprise multiple cell types with different treatment sensitivities, conventional therapy often fails to eradicate resistant subpopulations, which repopulate and drive relapse. Evolutionary therapy has therefore been proposed as a promising alternative strategy [29, 30]. Using principles from evolutionary game theory, evolutionary therapy aims to maintain a stable population of treatmentsensitive cells, which compete with resistant cells for limited space and resources and thereby suppress their expansion, rather than eliminating the sensitive population and leaving resistant cells free to proliferate unchecked. In neuroblastoma, how the three cell types interconvert in a specific patient is a key variable for this strategy, because interconversion can replenish a suppressed cell type from another, altering which subpopulation dominates and therefore which therapy is most effective over time. In this paper, we take an agnostic view on natural interconversion and assume that each of the 64 interconversion patterns between noradrenergic, mesenchymal, and I-type can be induced.

We built a mathematical model of three populations of neuroblastoma cells and used it to simulate how each interconversion pattern affects a phenotype-specific inhibitor’s ability to reduce heterogeneous neuroblastomas, including a sensitivity analysis on the drug dose and interconversion rate. This was followed by a stability analysis of the origin, which allowed us to quantify the effects of applying each interconversion pattern on remission maintenance. We identified interconversion patterns that effectively suppressed noradrenergic and mesenchymal cells respectively, and combined these with phenotype-specific inhibitors in multi-stage, cyclic treatment schedules to validate our findings in further numerical experiments.

## 2 Methods

Neuroblastoma (neuroblastoma) tumours contain three major cell populations which can interconvert and exhibit distinct treatment sensitivities. To investigate how interconversion patterns influence treatment efficacy, we built a model and simulations were conducted in three sequential phases: natural tumour growth (NTG), treatment, and relapse. In Natural tumour growth an equally small population of each cell type are allowed to grow without interference of an inhibitor. Treatment then adds either an N or S inhibitors causing drug dependant cell death. Finally, relapse has the inhibitor removed and the remaining populations after treatment are allowed to grow without inhibition.

### 2.1 Biological and Modelling Framework

The model will describe the interactions of the three types of neuroblastoma cells (N, I, S) in the presence of a single inhibitor applied continuously. The simulated drug targets either N or S cells, with potential off-target effects on the remaining cell types.

Each cell type undergoes three biological phenomena: logistic growth, drug induced cell death, and interconversion. These phenomena are described by a cell-type specific ordinary differential equation. There are 64 unique patterns of interconversion, arising from six possible directed pathways between the three cell types connecting edges, resulting in 2^6^ = 64 combinations.

The model was developed with state variables representing the population levels of N, S, and I, which are expressed as fractions of the carrying capacity. The carrying capacity was set at one.

## 2.2 Equations

The tumour population is defined as *P* and made up of the three population sizes *N, S* and *I*.

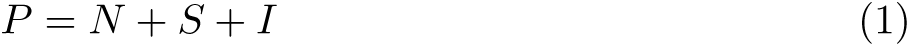

The model comprises three coupled non-linear first order ODEs, describing the population dynamics of the three cell types (Equations 2, 3, 4). Each equation includes:

1. A parameterized logistic growth term 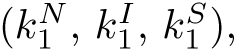,
2. a drug dependent death term 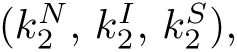,
3. intrinsic interconversion terms 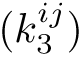 where *i* denotes the source cell type and *j* the target cell type.

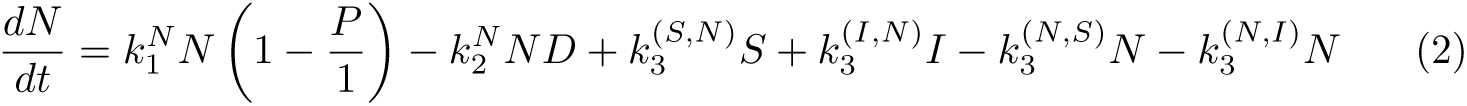

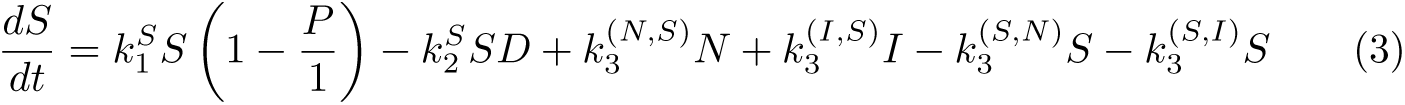

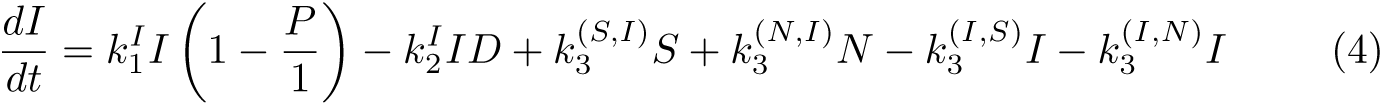

## 2.3 In Silico Drug Trial

A series of in-silico drug trials was performed to investigate the effects of inhibitors targeting N and S cell populations under the constraints of all 64 interconversion patterns. These simulations had three phases: Natural tumour growth(NTG), treatment and relapse.

Time *t* is expressed in units of days, such that an interval of *t* ∈ [0, 1] represents one day, based on reported neuroblastoma doubling times [11, 31–33]. The three biological phenomena — logistic growth, drug induced death, and interconversion – occur on the same time scale and are the same order of magnitude. The carrying capacity was taken to be a constant set to 1.

N cells were assigned the highest growth rate, S the lowest, and I an intermediate rate based on available biological evidence to mimic the three cell types seen in neuroblastoma. The non-target death rates, drug dose and interconversion rate were assigned as evenly spaced values in the absence of empirical rates. Because experimentally measured values are unavailable, non-target cell death rates were resistant (0) or susceptible (0.5), drug doses and interconversion rates were low (0.1), medium (0.5), and high (0.9) (Table 2).

Each experiment consists of three phases:

1. NTG, initial populations of all three cell types were set to 1 × 10*^−^*^6^ (Table 1). This represents a perturbation to the origin and an emerging tumour where no cell has yet established dominance. These initial populations were allowed to grow under the 64 interconversion pattern until the total population reached 0.4. This threshold reflects a tumour that has reached a detectable size.
2. For treatment phase a drug targeting either N or S cells was applied continuously in the interval of *t* ∈ [0, 100], reflecting typical 3–6 month clinical treatment durations [30, 34]. Additionally, this duration falls within the range reported in comparable pharmacokinetic modelling studies, which have used treatment periods of 150 days [35] and 208 days [36]. Death rates reflected known treatment sensitivities, and three drug doses were tested to capture clinically relevant variability (Table 2). Target cell death rate was set to 1 to represent maximal drug sensitivity (Table 1). The non-target cells were either unaffected or affected to a lesser degree than the target cell (Table 2), this allowed four unique plausible scenarios based on partial sensitivity or complete resistance of non-target cells. As exact values are unavailable, the tabulated values were selected to include both scenarios of partial and complete resistance. In the absence of exact empirical values, parameters were selected to capture these distinctions while remaining evenly spaced across a biologically reasonable range.
3. Relapse phase had the inhibitor removed and populations were allowed to grow again for the interval of *t* ∈ [0, 100] or until the total population reached 90%. This acts as a reasonable time frame of cell dynamics.

**Table 1:** Fixed parameter values used in the in-silico drug trial simulations. This table lists the baseline parameter values held constant across all simulations, including: initial cell population size (identical for all three cell types), intrinsic growth rates for each cell type (per time step), and drug induce death rate applied to the cell being pharmacologically targeted. These fixed parameters define the system’s core dynamical behaviour.

| Parameter | Value |
|---|---|
| Initial Cells | 0. 000001 |
| N Growth Rate (*k*<sub>1</sub>*N*) | 0.75 |
| I Growth Rate (*k*<sub>1</sub>*I*) | 0.5 |
| S Growth Rate (*k*<sub>1</sub>*S*) | 0.25 |
| Target Cell Death Rate (*k*<sub>2</sub>) | 1 |

**Table 2:** Parameter ranges explored in the in-silico drug trial simulations. This table specifies the parameter values varied across simulation runs: non target cell death rates for the two non-target cell types (0 or 0.5, giving four resistance/sensitivity combinations per pattern), drug doses (0.1, 0.5 or 0.9), and interconversion rates applied to each open paths in a pattern (low =0.1, medium = 0.5, or high = 0.9; closed paths are fixed at 0 by definition). These ranges were used to assess how treatment outcomes depend on drug strength and intrinsic interconversion dynamics affecting relative reduction scores and origin stability.

| Parameter | Value |
|---|---|
| Non-Target Cell Death Rate (*k*<sub>2</sub>) | {0, 0.5} |
| Drug Dose | {0.1, 0.5,0.9} |
| Interconversion Rates (*k*<sub>3</sub>) | {0, 0.1, 0.5, 0.9} |

A numerical experiment was performed for each pattern paired with each phenotypespecific inhibitor (64 ∗ 2 = 128 experiments). In each experiment, 3 ∗ 3 ∗ 4 = 36 runs of the three-phase sequence were conducted to cover every combination of 3 drug doses, 3 interconversion rates, and 4 resistance profiles (off-target death rates). Totalling of (36 ∗ 128) 4, 608 runs, this was done twice, once for each inhibitor type.

For each interconversion pattern, the relative reduction of the target population *X* ∈ {*N, S*} was calculated as the proportional decrease in treatment and relapse phases. In each experiment, we calculated 36 relative reduction values for the treatment phase and 36 for the relapse phase, corresponding to the 36 parameter combinations tested. For each phase, we counted the number of parameter combinations that yielded a non-negative relative reduction to obtain a score, Score*_p_*, out of 36.

Interconversion patterns were shortlisted based on the following procedures.

1. If N or S population at the end of NTG *<* 0.05, the pattern was selected for the next step.
2. These patterns were then ordered by the lowest mean final I population, and patterns with isolated I cells (“|I|”) were removed from consideration. This filtering step is biologically motivated: I represent I-type cells, and their presence is associated with poor clinical outcomes [3, 15, 20]. Patterns that permit the unchecked accumulation of I-type cells are therefore considered clinically undesirable, regardless of their effect on the target population.
3. If the N population *<* 0.05 at the end of NTG, the pattern was classified as N-suppressing and an S inhibitor was applied during treatment. Vice versa was done for S-suppressing patterns. The top 2 N-suppressing and top 3 S-suppressing patterns were selected for further analysis.
4. *Score_X,T_* (out of 36) and *Score_X,R_*(out of 36) were calculated for the cell population targeted by the inhibitor.

## 2.4 Sensitivity Analysis

Five interconversion patterns of interest were identified during the in silico drug trial (Table 7). Two were N-suppressing and three were S-suppressing. For each cell type the population at the end of NTG was averaged across the three drug doses and four non-target cell death rate combinations for each pattern.

During the treatment phase, the inhibitor that synergised with the pattern was used, with the inhibitor suppressing the opposite cell to the pattern. N-suppressing patterns had an S inhibitor applied. S-suppressing patterns had an N inhibitor applied. For each experiment and each of the three drug doses, *Score_T_* and *Score_R_* were calculated using the results of the 12 corresponding runs. These procedures resulted in six dose-specific scores for each experiment, including three for each phase.

The same sensitivity analysis was carried out dividing each set if 36 runs into three subsets for the interconversion rate, following the same procedure described above, with *Score_X,T_*, *Score_X,R_* evaluated across low, medium and high interconversion rates.

## 2.5 Stability Analysis

To understand how remission could be maintained by inducing a shortlisted pattern and its synergistic inhibitor, a stability analysis was performed for the origin in each of the five cases. For each of these combinations, interconversion rates and drug doses between ∈ [0.1 − 0.9] were considered for the four combinations of non-target cell death rates (Table 3). When relevant, the drug dose was held constant. Here, we define remission as a stable origin during treatment. The Jacobian matrix at the origin was used to compute eigenvalues, and stability was determined by the sign of the largest real part.

**Table 3:** Parameter values used for numerical stability analysis. The Jacobian was evaluated at the origin (initial guesses = 0 for all cell types), distinct from the small NTG initial population used in the in-silico drug trial. The growth rates for each cell type, target and non-target cell death rates match Table 1. Drug dose and interconversion rate waried from 0.1 to 0.9. the five interconversion patterns of interest (P1-P5) are listed using the bracket notation described in Figure 1.

| Parameter | Value |
|---|---|
| Initial Guesses | 0 |
| N Growth Rate (*k*<sub>1</sub>*N*) | 0.75 |
| I Growth Rate (*k*<sub>1</sub>*I*) | 0.5 |
| S Growth Rate (*k*<sub>1</sub>*S*) | 0.25 |
| Target Cell Death Rate (*k*<sub>2</sub>) | 1 |
| Non-Target Cell Death Rate (*k*<sub>2</sub>) | {0, 0.5} |
| Drug Dose | 0.1, 0.2, 0.3, 0.4, 0.5, 0.6, 0.7, 0.8, 0.9 |
| Interconversion Rates (*k*<sub>3</sub>) | 0.1, 0.2, 0.3, 0.4, 0.5, 0.6, 0.7, 0.8, 0.9 |
| Interconversion patterns | {<N> \|I> \|S\|, <N> \|I><S\|, \|N\| <I> \|S>, \|N\| <I> \|S\|, \|N\| <I><S>} |

Results were grouped by non-target cell death rate combination, with four scenarios of partial or complete resistance of non-target cells to the inhibitor. Heatmaps of the maximum real eigenvalue were generated across drug dose and interconversion rate to examine the dependence of stability on these parameters.

## 2.6 Simulating Hypothetical Treatment Schedules for Validation

Based on the previous analysis, P1, P3 and P4 were selected as the medically useful patterns, to examine the effects of different treatment schedules. P2 scored lower in the sensitivity analysis and stability analysis than P1. Additionally, the literature indicates that P2 and P5 are unlikely to occur in an untreated neuroblastoma[18]. Therefore, they were removed from the shortlist. To validate the final shortlist, we proposed the following treatment schedules combining P1, P3, and P4 with the two phenotype-specific inhibitors.

We carried out simulations where hypothetical treatment schedules were applied by cycling between a sequence of phases in each simulation. Each treatment schedule was tested on 200 virtual tumours with different initial compositions. These initial conditions included: 50 N-dominant with N cells making up *>* 50% of the population, 50 I-dominant I cells making up *>* 50% of the population, 50 S-dominant S cells making up *>* 50% of the population, and 50 compositions with no dominant cell type. Four treatment schedules were tested, each was repeated for four cycles.

The two-phase schedule did not involve the use of an inhibitor (Table 4 A). Either P3 or P4 was applied in phase one, followed by P1 in phase two. The three-phase treatment schedule builds on this with an extra phase in the middle. In this new phase, the inhibitor that complements with the pattern used in the first phase is used alongside the pattern (P3/P4 plus N inhibitor or P1 plus S inhibitor) (Table 4 B and C).

**Table 4:** Overview of the four treatment schedules used in the validation analysis. Each row shows the pattern ID and where applicable inhibitor where applied in each phase. Schedules with fewer than four phases leave later columns blank.

| Schedule | Phase 1 | Phase 2 | Phase 3 | Phase 4 |
|---|---|---|---|---|
| A (two-phase) | P3 or P4 | P1 | – | – |
| B (three-phase) | P1 | P1 + S inhibitor | P3 or P4 | – |
| C (three-phase) | P3 or P4 | P3 or P4 + N inhibitor | P1 | – |
| D (four-phase) | P3 or P4 | P3 or P4 + N inhibitor | P1 | P1 + S inhibitor |

The four-phase schedules (Table 4 D), phase one applied S-suppressing P3 or 4 with no inhibitor, then N inhibitor was added to the same pattern in phase two. In phase three the N inhibitor was removed and P1 applied, followed phase four by S inhibitor deing applied to P1.

Phases one and three were run for 30 time steps and Phases two and four combining inhibitor and pattern ran for 100 time steps. Each treatment schedule was applied to 200 initial conditions twice, once with P3 and once with P4 in phase one. For each set of initial conditions and each treatment schedule the drug dose was set to 0.9 for phases with an active inhibitor, the interconversion rate was set to 0.5, and four combinations of death rates were used for the non-target cell types.

## 3 Results

### 3.1 in-silico Drug Trial

The in silico drug trial results indicate that six of the 64 patterns led to a mean population of N below 0.05 at the end of NTG (Table 5). Thirty-three of the patterns achieved the same with respect to the population of S (Table 6). Patterns with isolated I cells (“|I|”) were removed from consideration as I-type cells are associated with poor clinical outcomes [21].

**Table 5:** Mean N, I, and S populations across interconversion patterns leading to mean N population ¡0.05.

| **Interconversion Pattern** | **Mean N** | **Mean I** | **Mean S** |
|---|---|---|---|
| <N> \|I><S\| | 0.034995447 | 0.163494771 | 0.128489178 |
| <N> \|I> \|S\| | 0.049628901 | 0.04003377 | 0.291466801 |
| <N> \|I\| <S> | 0.034725108 | 0.285871501 | 0.007859875 |
| <N> \|I\| <S\| | 0.031318952 | 0.281777124 | 0.00979586 |
| <N> \|I\|\|S> | 0.038826664 | 0.266909541 | 0.011024758 |
| <N> \|I\|\|S\| | 0.032759447 | 0.260588702 | 0.014350547 |

**Table 6:** Mean N, I, and S populations across interconversion patterns leading to mean S population ¡0.05.

| Interconversion Pattern | Mean N | Mean I | Mean S |
|---|---|---|---|
| \|N\| <I> \|S> | 0.27825082 | 4.24284 × 10<sup>−5</sup> | 3.28968 × 10<sup>−5</sup> |
| \|N\| <I> \|S\| | 0.307308461 | 4.39401 × 10<sup>−5</sup> | 0.000143096 |
| \|N\| <I><S> | 0.283941058 | 9.33784 × 10<sup>−5</sup> | 3.56241 × 10<sup>−5</sup> |
| \|N\| <I><S\| | 0.298428352 | 0.000125615 | 7.10965 × 10<sup>−5</sup> |
| \|N\| <I\|\|S> | 0.280173383 | 0.000232156 | 3.76419 × 10<sup>−6</sup> |
| \|N\| <I\|\|S\| | 0.336129798 | 0.000238076 | 6.11693 × 10<sup>−5</sup> |
| \|N\|\|I> \|S> | 0.299795586 | 0.000240611 | 0.000107011 |
| \|N\|\|I> \|S\| | 0.240704372 | 0.000248928 | 0.000300071 |
| \|N\| <I\| <S> | 0.287830812 | 0.000297719 | 5.68182 × 10<sup>−7</sup> |
| \|N\| <I\| <S\| | 0.316965905 | 0.000329913 | 3.79671 × 10<sup>−6</sup> |
| \|N\|\|I><S> | 0.294075456 | 0.000471132 | 0.000139529 |
| \|N\|\|I><S\| | 0.243723789 | 0.000986425 | 0.000580368 |
| \|N\|\|I\|\|S> | 0.313304819 | 0.003788529 | 3.84055 × 10<sup>−6</sup> |
| \|N\|\|I\|\|S\| | 0.238998733 | 0.004046263 | 6.38646 × 10<sup>−5</sup> |
| \|N\|\|I\| <S> | 0.295634379 | 0.005207682 | 8.10672 × 10<sup>−7</sup> |
| \|N\|\|I\| <S\| | 0.239336748 | 0.006371128 | 3.89059 × 10<sup>−6</sup> |
| \|N><I> \|S> | 0.19681809 | 0.081205751 | 0.049309646 |
| \|N><I><S> | 0.193359543 | 0.094006441 | 0.034211182 |
| \|N><I\| <S\| | 0.174274697 | 0.119083327 | 4.43177 × 10<sup>−6</sup> |
| \|N><I\|\|S\| | 0.177250093 | 0.123633743 | 0.000102005 |
| \|N><I\|\|S> | 0.181735485 | 0.123686648 | 4.39526 × 10<sup>−6</sup> |
| \|N><I\| <S> | 0.185899701 | 0.127015661 | 7.29297 × 10<sup>−7</sup> |
| <N\|\|I\| <S\| | 0.080855006 | 0.22782943 | 0.019555759 |
| <N\|\|I\|\|S\| | 0.083061284 | 0.231858595 | 0.028447794 |
| <N\|\|I\| <S> | 0.08193339 | 0.239746603 | 0.017486448 |
| \|N> \|I\|\|S> | 0.057884841 | 0.259151183 | 5.01294 × 10<sup>−6</sup> |
| \|N> \|I\| <S> | 0.056588623 | 0.259337644 | 7.60546 × 10<sup>−7</sup> |
| <N> \|I\|\|S\| | 0.032759447 | 0.260588702 | 0.014350547 |
| <N> \|I\|\|S> | 0.038826664 | 0.266909541 | 0.011024758 |
| \|N> \|I\|\|S\| | 0.050813827 | 0.269092728 | 0.000281907 |
| \|N> \|I\| <S\| | 0.05063688 | 0.271860735 | 5.10445 × 10<sup>−6</sup> |
| <N> \|I\| <S\| | 0.031318952 | 0.281777124 | 0.00979586 |
| <N> \|I\| <S> | 0.034725108 | 0.285871501 | 0.007859875 |

After the second filtering step, two N-suppressing patterns and 18 S-suppressing patterns remained. From the 18 S patterns, the three patterns with highest scores based on relative reduction were selected as S-suppressing (“|N| |S*>*”, “|N| |S|”, “|N| *<*S*>*”) and used along side the patterns for N (“*<*N*>* |I*><*S|”, “*<*N*>* |I*>* |S|”) for detailed analysis (Table 7). Here, ‘*<*’ ‘*>*’ denotes an open path and direction of interconversion, and ‘|’ denotes a closed path of a cell type, as defined in Section 2.2.

**Table 7:** Patterns of interest selected for downstream analysis. interconversion pattern between noradrenergic like (N), intermediate (I), and Mesenchymal like (S) cell types are shown for five pattens shortlisted in Section 2.2 (P1-P2, N-suppressing; P3-P5, S-suppressing). Using the notation defined in Figure 1 and Blue arrows (→) indicate positive flow.

| Target Cell | Pattern |  |  |
|---|---|---|---|
| N-suppressing | P1:<N> \|I> \|S\| |  | P2:<N> \|I><S\| |
| S-suppressing | P3:\|N\| <I> \|S> | P4:\|N\| <I> \|S\| | P5:\|N\| <I><S> |

Complementary inhibitors were applied to these patterns; N-suppressing patterns were treated with the S inhibitor and vice versa.

The S-suppressing patterns (P3-5) were more effective against S cells than the Nsuppressing patterns (P1 and P2) were against N cells during NTG. Additionally, all three S-suppressing patterns maintained final mean I populations at *<* 1 × 10*^−^*^3^ during NTG. N cell are the fastest growing which explains the discrepancy in the pattern’s ability to suppress S or N cells.

When an inhibitor of the unsuppressed cell types is applied, the S inhibitor is applied to N-suppressing patterns and vice versa; the S-suppressing patterns express higher relative reduction scores during treatment and maintain higher scores during relapse.

Figure 2 shows two representative examples of the in silico drug trial output, illustrating how a pattern and its complementary inhibitor interact across the three phases of a simulation. Figure 2a shows a time-series plot for the N-suppressing pattern P1 under a medium interconversion rate and a high S inhibitor dose. The P1 time series plot shows that S is the dominant cell at the end of NTG, with N and I approximately zero. The S inhibitor can suppress the remaining cells during treantment. In contrast, Figure 2b shows the S-suppressing pattern P3 with medium interconversion rate, under a high N inhibitor dose. At the end of NTG N cells are present and are suppressed by the N inhibitor during treatment. However, N can re-emerge during relapse when the N inhibitor is removed.

**Fig. 1:**
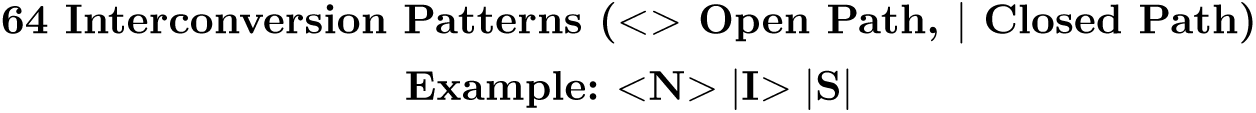
Notation for the 64 interconversion patterns. Each cell type is written with two brackets, one on each side, describing whether that cell type’s own outgoing interconversion path to each neighbour is open or closed. Angle brackets (*<* or*>*) denote an open path between cell types in that direction. A vertical bar (|) denotes a closed path (no interconversion flow between adjacent cell types). The example shown, *<*N*>* |I*>* |S|, indicates that N can convert bidirectionally into both I and S, I can only convert into S, and S can convert into neither I nor N.

**Fig. 2:**
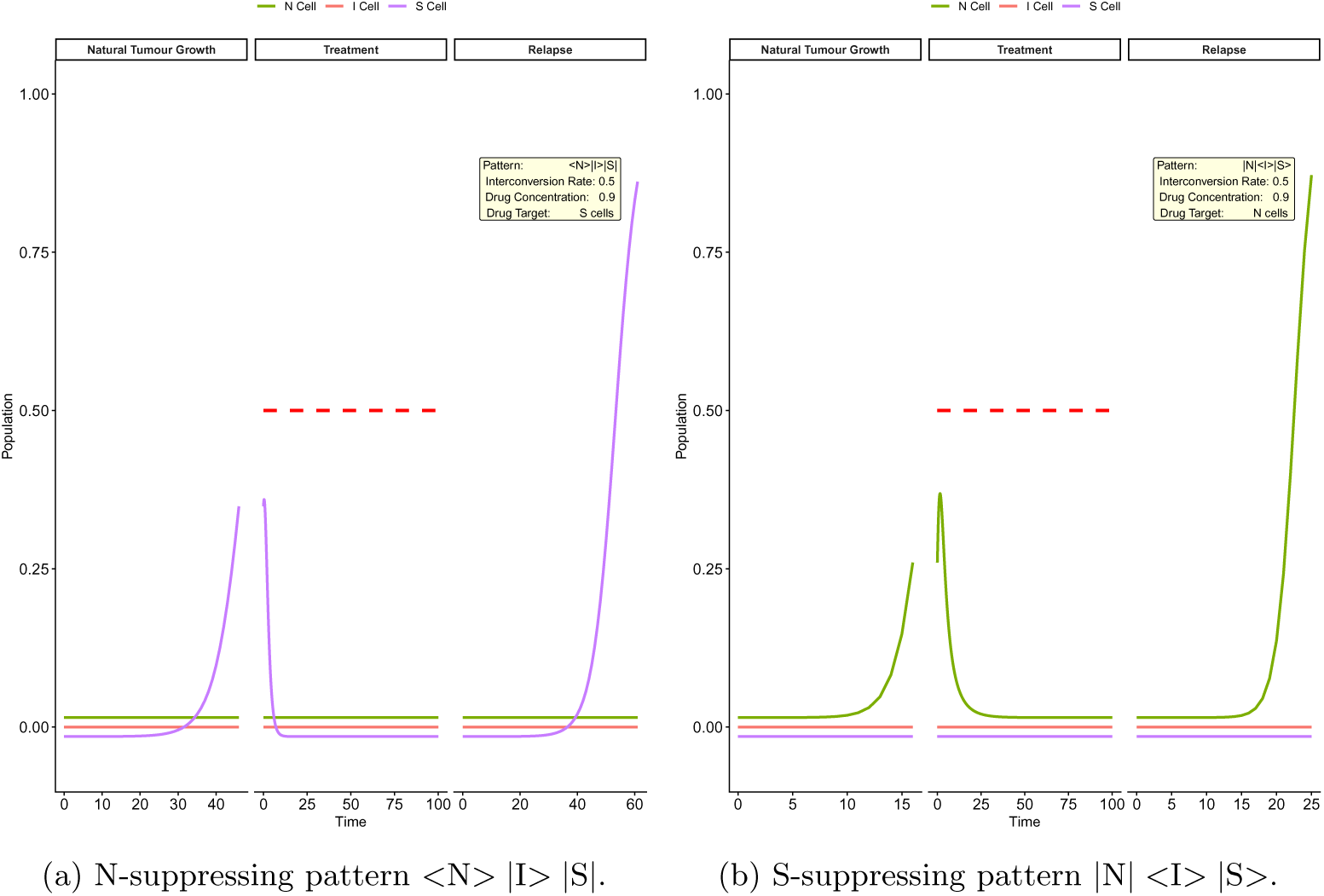
Representative examples of in silico drug trial output, illustrating time series of natural tumour growth, treatment, and relapse phases for one N-suppressing and one S-suppressing pattern. Cell population (fraction of carrying capacity) is shown on the y-axis against time in days on the x-axis. N cells (green), I cells (pink), and S cells (purple) are shown alongside drug concentration (dashed red). For visual clarity, overlapping populations near zero are offset slightly, population values are otherwise unmodified. (a)N-suppressing pattern *<*N*>* |I*>* |S| under S inhibitor (interconversion rate = 0.5, drug concentration = 0.9). During natural tumour growth only S cells are present. S cells are rapidly suppressed to zero during treatment and do not re-emerge during relapse. (b)S-suppressing pattern |N| |S*>* under N inhibitor (interconversion rate = 0.5, drug concentration = 0.9). All three cell types are present during natural tumour growth. N and I cells are suppressed during treatment but re-emerge during relapse, approaching carrying capacity.

This indicates that, with the appropriate inhibitor, P1 populations reach an extinct state during treatment and take longer to relapse, once the inhibitor is withdrawn. In terms of the underlying dynamical system, this slower relapse is consistent with this pattern-inhibitor combination lying close to a stable region of parameter space. The origin of P3 under an N inhibitor is less stable in this example, allowing N to reemerge more quickly once the inhibitor is removed. Here we classify scenarios where population extinction from treatment is stable as remission, and scenarios where cells can re-emerge as relapse. Further stability analysis was carried out after analysing parametric sensitivity to find true stable parameter combinations.

### 3.2 Sensitivity Analysis

Figure 3 shows, for each of the five shortlisted patterns, how sensitive its suppression of each cell type is to the interconversion rate during NTG. Each row is a pattern, and each column shows the mean cell populations at low, medium, and high interconversion rates. This allows us to check whether a pattern’s ability to suppress cell types depends on how fast cells interconvert, before any inhibitor is applied. The N-suppressing patterns permit interconversion from N into both I and S cell types (Table 8). The results suggest that P1 and P2 require at least a medium interconversion rate to work effectively against N cells. Furthermore, P1 can suppress I cells with a higher interconversion rate. In short, Figure 3 indicates that the three S-suppressing patterns are effective against S and I cells during NTG regardless of how high the interconversion rate is, whilst the two N-suppressing patterns are more sensitive to this parameter during NTG.

**Fig. 3:**
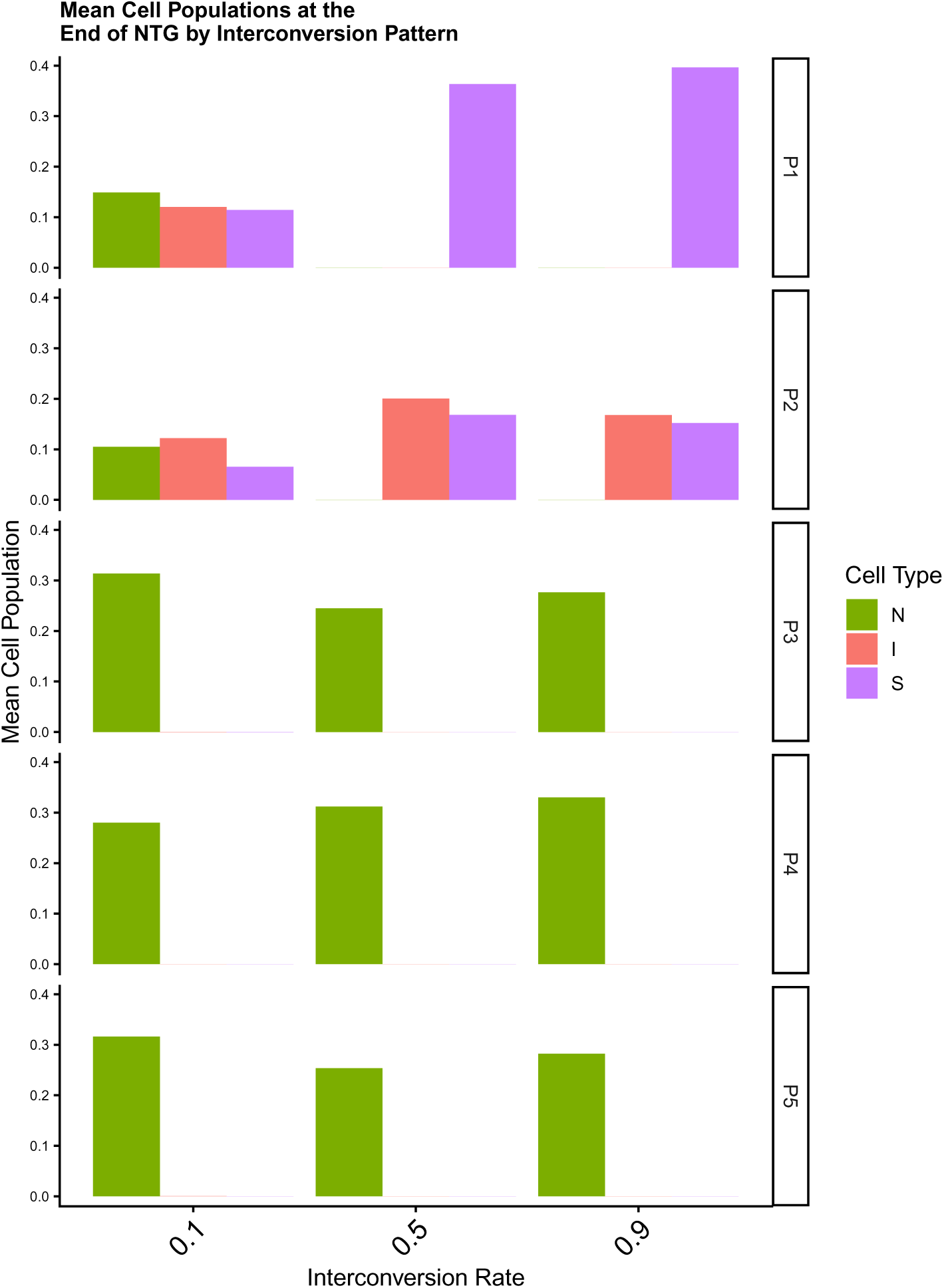
Mean cell populations (coloured by cell type) at the end of the natural tumour growth phase, faceted by pattern and interconversion rate. Each row shows one of the five patterns of interest: the top two rows show N-suppressing patterns and the bottom three rows show S-suppressing patterns. Columns correspond to low, medium, and high interconversion rates.

**Table 8:** Summary of NTG mean cell populations and cross-inhibitor performance scores for patterns of interest. N-suppressing pattern (P1, P2) were treated with S inhibitor. S-suppressing patterns (P3-P5) were treated with N inhibitor. T Score and R Score are based on the relative reduction during treatment and relapse phases for number of parameter combinations (out of 36).

| Pattern ID | Interconversion Pattern | NTG<br>Mean N | NTG<br>Mean I | NTG<br>Mean S | T<br>Score | R<br>Score |
|---|---|---|---|---|---|---|
| P1 | <N> \|I> \|S\| | 0.050 | 0.040 | 0.291 | 9 | 0 |
| P2 | <N> \|I><S\| | 0.035 | 0.163 | 0.128 | 4 | 7 |
| P3 | \|N\| <I> \|S> | 0.278 | 4.24 × 10<sup>−5</sup> | 3.29 × 10<sup>−5</sup> | 15 | 11 |
| P4 | \|N\| <I> \|S\| | 0.307 | 4.39 × 10<sup>−5</sup> | 1.43 × 10<sup>−4</sup> | 19 | 11 |
| P5 | \|N\| <I><S> | 0.284 | 9.34 × 10<sup>−5</sup> | 3.56 × 10<sup>−5</sup> | 14 | 11 |

Complementary inhibitors were used during treatment: N-suppressing patterns had the S inhibitor applied, and S-suppressing patterns had the N inhibitor applied. As a result, the relative reduction scores of the inhibitor-targeted cells were assessed. The number of runs leading to a positive relative reduction was counted to get relative reduction scores.

#### 3.2.1 Drug Concentration

Figure 4 illustrates how the five patterns enhance their respective complementary inhibitors at different doses. As a reminder, P1 and P2 are effective against N cells, so their complementary inhibitor is effective against S cells. The results indicate that P1 combines with the inhibitor more effectively, achieving better reduction scores at the same doses. P1 achieved S reduction for more of the parameter space at medium or high drug doses compared to P2 (treatment score 9/12 and 12/12 versus 4/12 and 9/12 for P2)(Figure 4a). Conversely, P1 was unable to reduce or maintain a population of S close to zero during relapse.

**Fig. 4:**
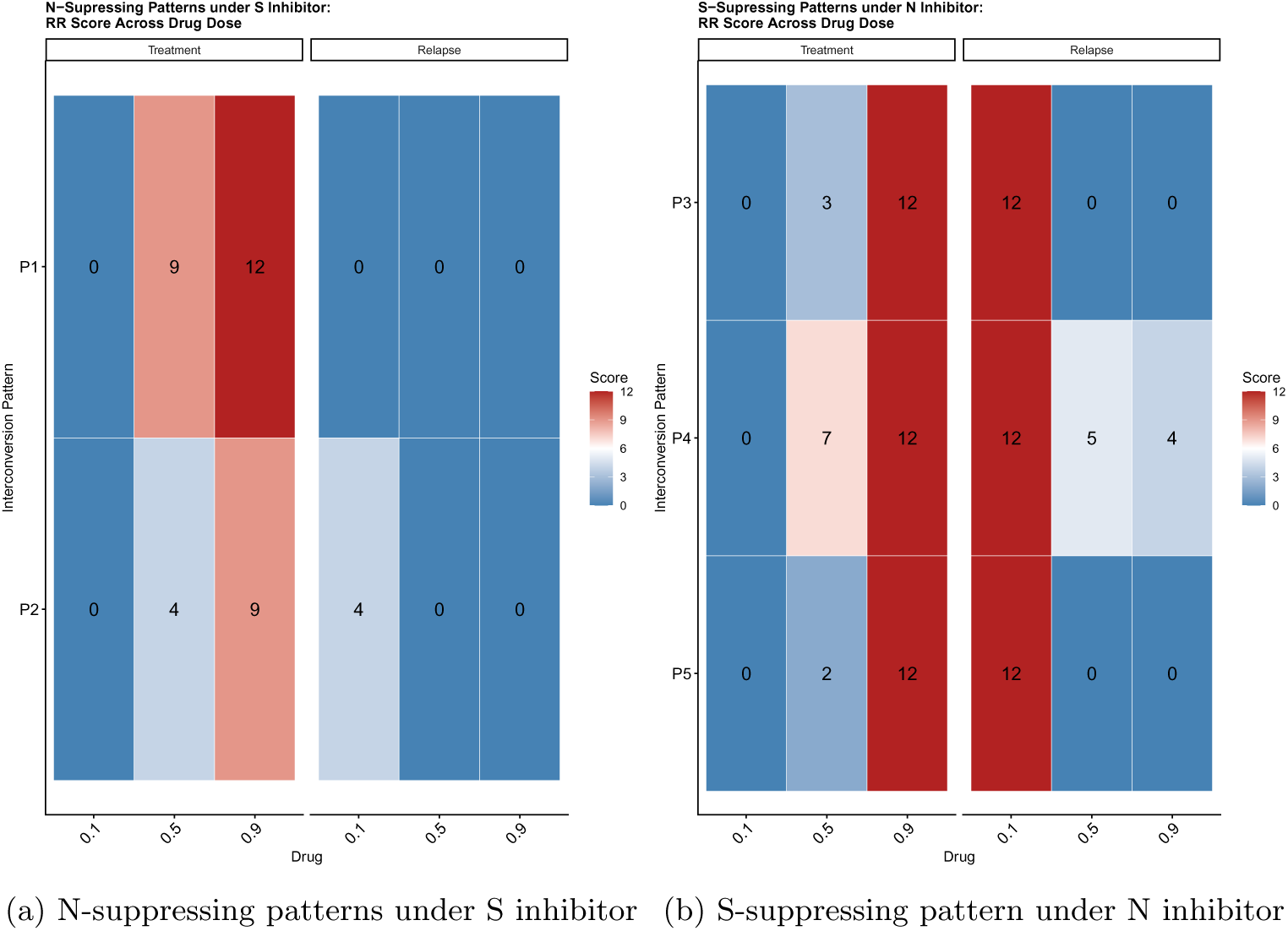
T Score and R Score from Table 8 (out of 36 total combinations), broken down by drug dose: for each dose (low = 0.1, medium = 0.5, high = 0.9), the score out of 12 sums over all three interconversion rates and all non-target cell death-rates. (a) Nsuppressing patterns (P1, P2), scored for reduction of the S population under the S inhibitor, during treatment and relapse phases. (b) S-suppressing patterns (P3–P5), scored for reduction of the N population under the N inhibitor, shown during treatment and relapse phases. Colour scale: blue = low score, red = high score.

P4 (|N| |S|) is better than the other two when a medium dose is applied. This is because S is the slowest-growing cell and acts as a sink for I cells and cannot become N. However, this will lead to an S-dominant population which would be detrimental to the patient if not treated with an S targeting therapy.

#### 3.2.2 Interconversion Rate

Figure 5 presents equivalent results to Figure 4 for the interconversion rate. Nsuppressing patterns treated with an S inhibitor were scored for S-relative reduction. During treatment, P1 had the highest scores over all patterns but was most successful on medium-to-high interconversion rates (Figure 5a). This is because S is isolated and becomes a sink for N and I; therefore, higher interconversion rates make N and I into S and are reduced faster by the S inhibitor, in addition to S having the lowest proliferation rate.

**Fig. 5:**
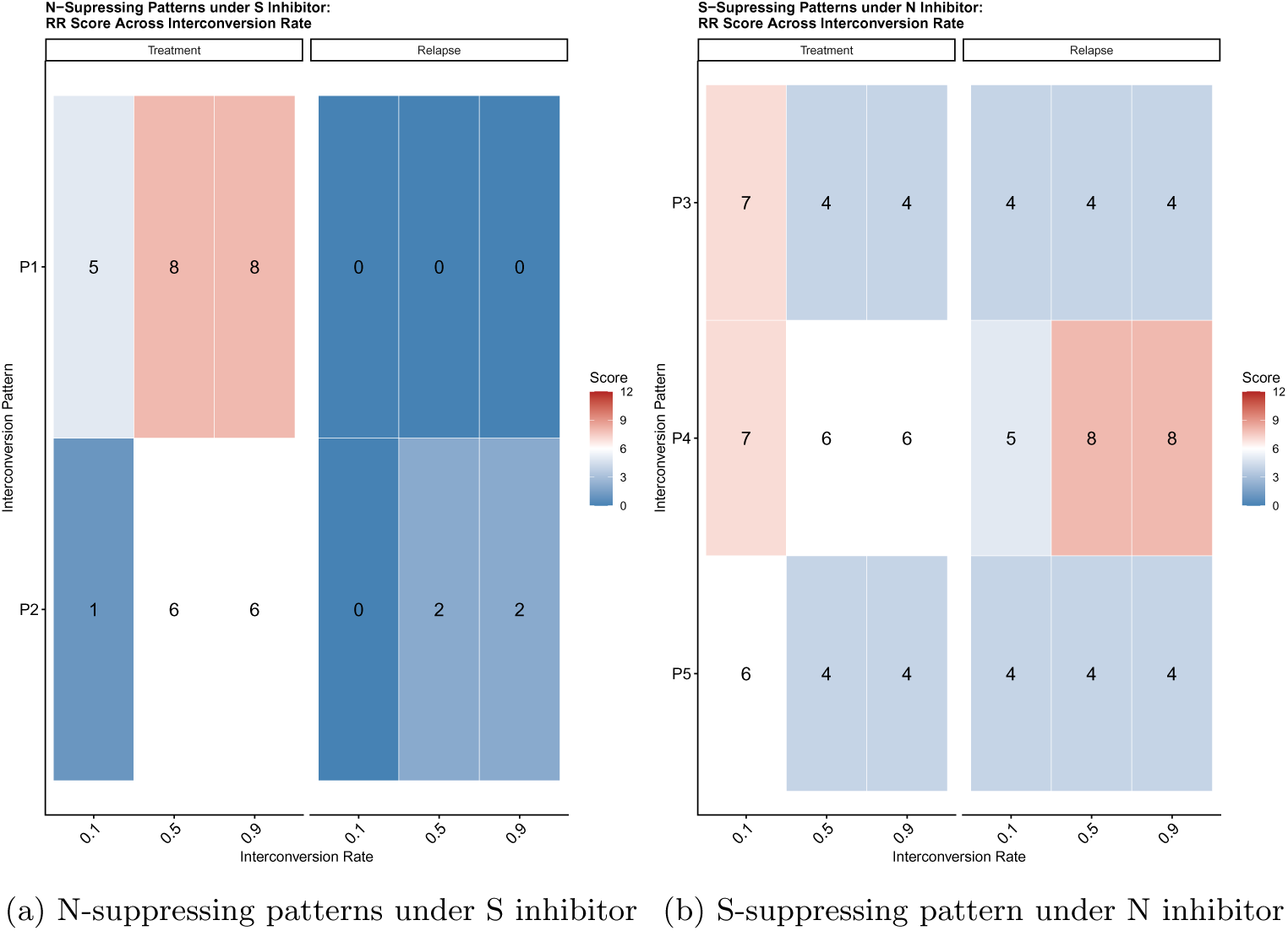
T Score and R Score from Table 8 (out of 36 total combinations), broken down by interconversion rate: for each rate (low = 0.1, medium = 0.5, high = 0.9), the socore out of 12 sums over all three drug doses and non-target cell death-rates. (a) N-suppressing patterns (P1, P2), scored for reduction of the S population under the S inhibitor, during treatment and relapse phases. (b) S-suppressing patterns (P3–P5), scored for reduction of the N population under the N inhibitor, shown during treatment and relapse phases. Colour scale: blue = low score, red = high score.

The results suggest that all three S-suppressing patterns enhance an N inhibitor most effectively when interconversion is slow. This is due to N being isolated in these patterns and being the fastest growing between the three cell types. Reducing the influx into N gives the N inhibitor more time to eliminate them before they can grow exponentially.

P4 also maintained higher scores at medium and high interconversion rates during treatment (Figure 5b). Combined with its performance across drug doses, P4 appears less sensitive to variation in both drug concentration and interconversion rate. This may be attributed to the dual isolation of N and S in P4: interconversion is directed toward S, which grows slowly and poses less clinical risk, or toward N, which is targeted directly by the inhibitor. This suggests that P4 was the best pattern for reducing or maintaining N populations, especially at low interconversion rates. However, this will lead to an S-dominant population which would be detrimental to the patient as S represents MES which are more resistant to treatment and more migratory.

### 3.3 Stability Analysis

The Jacobian matrix of the system (Equations 5), derived by linearising the system of three ODEs (Equations 2-4), was evaluated at the origin to assess local stability. Where the Jacobian evaluated at the steady state is:

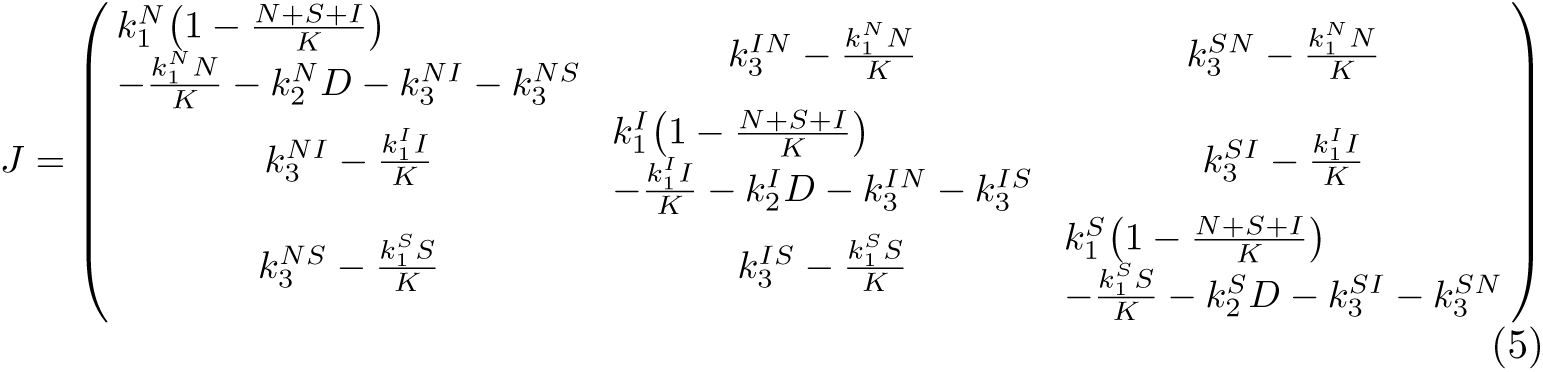

For each parameter combination, the eigenvalues were computed. The eigenvalue with the largest real part determines whether the origin is stable at the parametric combination. A negative value indicates stability, meaning the origin is an attracting fixed point and the system’ state vector will return to it following a small perturbation, while a positive value indicates instability. This analysis was csarried out for P1, P2, P3, P4, and P5. For each pattern, four groups of parametric combinations corresponding to the four combinations of non-target death rates were considered. In each group, 10 death rates (including 0) and nine interconversion rates were considered.

#### 3.3.1 Characterisation of the Origin of N-suppressing patterns under S inhibitor

Across P1 and P2, we found that stabilising the origin required a sufficiently high drug dose, and both patterns benefited from higher interconversion rates. P1 could be stabilised with a lower drug dose than P2 at every non-target death rate combination tested, reflecting the I-type reservoir in P2 leading to drug escape. The results for each non-target death rate combination are detailed below.

The stability of the origin was assessed for P1 and P2 across all four combinations of non-target death rates (Figure 6). The origin is always unstable when the drug dose is zero, as expected due to the logistic growth of the system. At baseline (*K*2*_N_* = 0, *K*2*_I_* = 0), P1 exhibited a bifurcation with respect to two parameters: above an interconversion rate of 0.5, drug doses greater than 0.5 were sufficient to stabilise the system (Figure 6a). The maximum real part correlates negatively with both parameters. P2 remained entirely unstable across all drug doses and interconversion rates at baseline, consistent with the I-cell reservoir enabling drug escape.

**Fig. 6:**
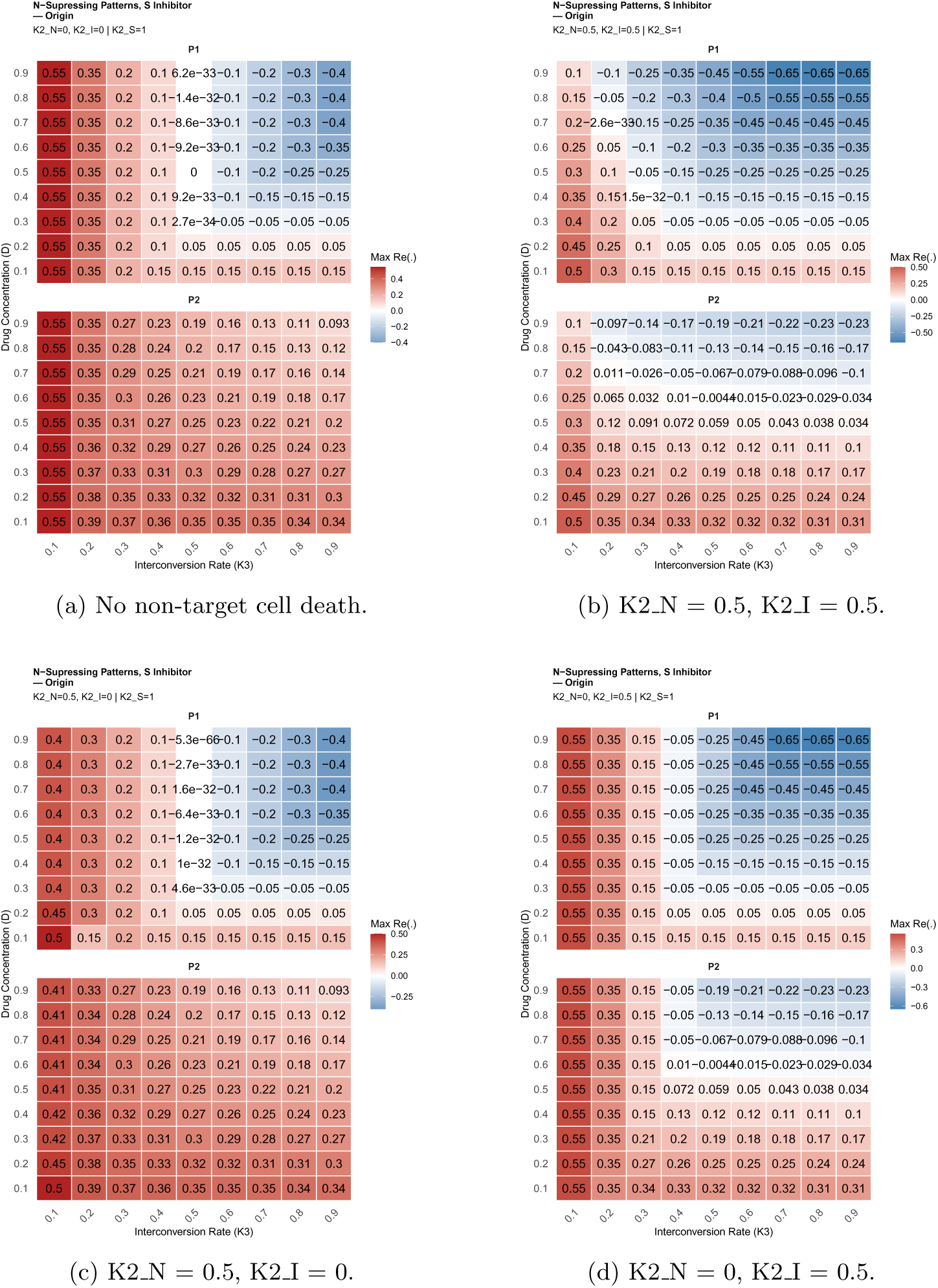
Stability of N-suppressing patterns under S inhibitor across all non-target cell death rate combinations. Each subfigure shows a heatmap of the maximum real part of the eigenvalue at the origin, with blue indicating stability and red indicating instability. K2 S = 1 throughout.

When both non-target death rates were set to 0.5, a further improvement was observed – the bifurcation point shifted to an interconversion rate of 0.2, with minimum drug doses of *D* ≥ 0.3 for P1 and *D* ≥ 0.6 for P2 (Figure 6b). Setting *K*2*_N_* = 0.5 alone produced no change from baseline in either pattern (Figure 6c), confirming that N-cell death alone is insufficient for stabilisation. In contrast, setting *K*2*_I_* = 0.5 alone improved stability for both patterns (Figure 6d), shifting the bifurcation point to an interconversion rate of 0.4 for both P1 and P2, with minimum drug doses of *D* ≥ 0.3 and *D* ≥ 0.6, respectively. Taken together, drug dose and interconversion rate are the primary drivers of stability for N-suppressing patterns under the S inhibitor, with *K*2*_I_* impacting their efficacy attributable to its role in eliminating the I-cell reservoir, with I-type cells enabling drug escape.

#### 3.3.2 Origin of S-suppressing patterns under N inhibitor

Across P3, P4, and P5, we found that we can generally stabilise the origin with a sufficiently high N inhibitor dose, and that the required drug dose and interconversion rate depend on whether non-target cell death is present. P3 and P5 can be stabilised by drug dose and interconversion rate alone, without any non-target cell death. P4 cannot, because S cells have no outgoing interconversion path in this pattern; S accumulates unchecked, and the origin remains unstable regardless of drug dose or interconversion rate unless non-target cell death also clears this accumulation. When non-target cell death rates for both S and I increase, all three patterns become stabilisable, and drug dose alone then determines the outcome. The results for each non-target death rate combination are detailed below.

The stability of the origin was assessed for P3, P4, and P5 across all four non-target death rate combinations (Figure 7). At baseline (*K*2*_S_* = 0, *K*2*_I_* = 0), P3 achieved stability at an interconversion rate of *K*3 ≥ 0.3, and P5 at *K*3 ≥ 0.4, both with *D*≥ 0.8 (Figure 7a). Once the bifurcation threshold was crossed for P3, the interconversion rate had no further effect on the stability depth — the drug dose was the sole determinant. P4 remained entirely unstable at baseline, with the lowest observed eigenvalue of 0.25 corresponding to the S growth rate, suggesting S accumulation as the destabilising mechanism.

**Fig. 7:**
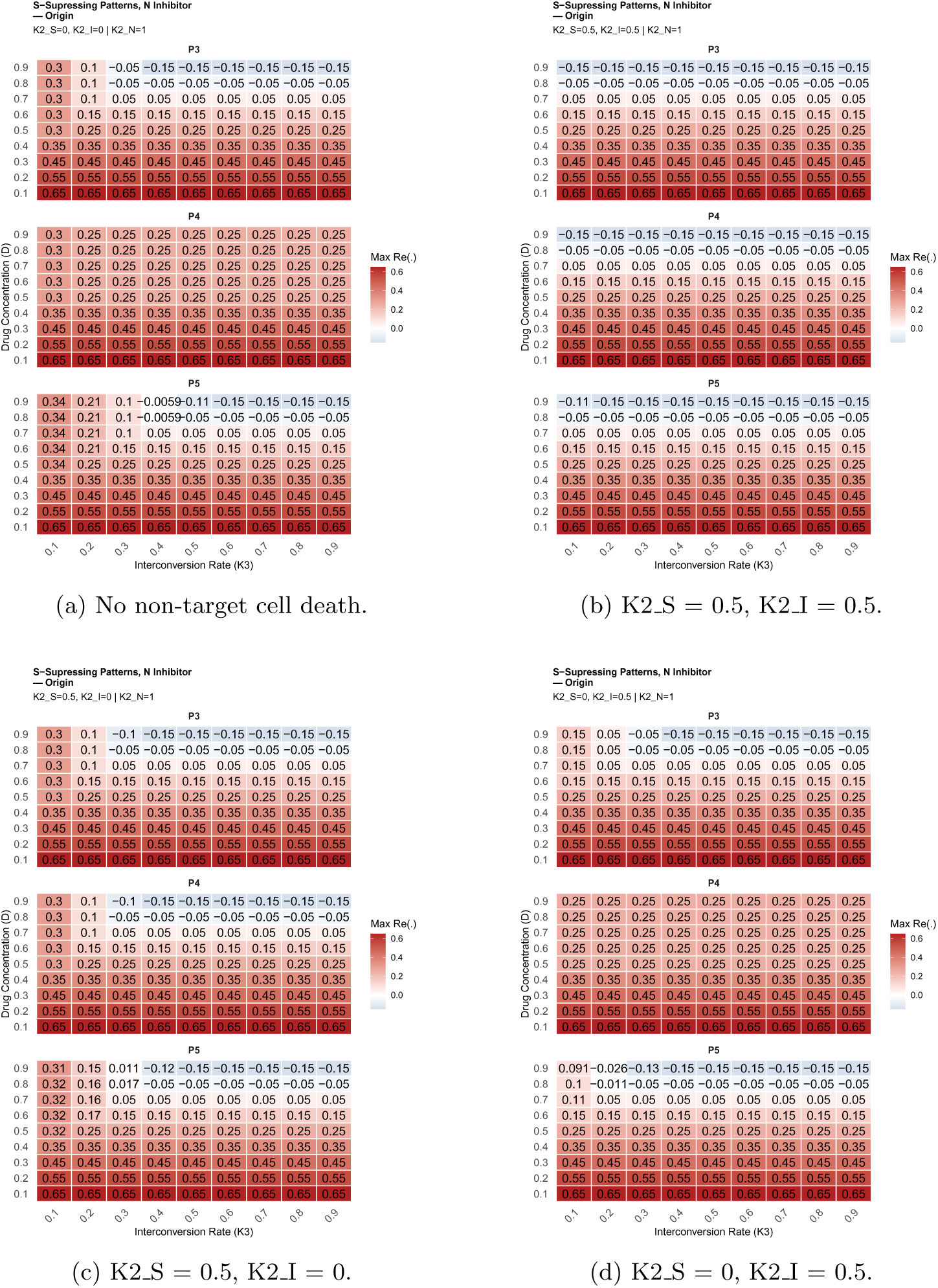
Stability of S-suppressing patterns under N inhibitor across all non-target cell death rate combinations. Each subfigure shows a heatmap of the maximum real part of the eigenvalue at the origin, with blue indicating stability and red indicating instability. K2 S = 1 throughout.

Further supporting this idea (S accumulation is destabilising), setting *K*2*_S_* = 0.5 alone stabilised P3 and P4 at *K*3 ≥ 0.3 and P5 at *K*3 ≥ 0.4, all requiring *D >* 0.8 (Figure 7c). The effect of *K*2*_S_* is likely driven by I cells transitioning into the slowgrowing S population, which is then further suppressed by S death, preventing escape into faster-growing states. Setting *K*2*_I_* = 0.5 alone maintained P3 at *K*3 ≥ 0.3 but improved P5 to *K*3 ≥ 0.2, suggesting *K*2*_I_* has a stronger effect on P5 than *K*2*_S_* (Figure 7d). This follows the pattern seen at baseline (*K*2*_S_* = 0, *K*2*_I_* = 0), with the origin’s stability being unaffected by the interconversion rate. However, P4 remained unstable when *K*2*_I_* alone was set to 0.5. In P4, S is isolated with no outgoing flow and therefore acts as a terminal sink; without *K*2*_S_*, S cells accumulate unchecked, and the system cannot be stabilised. When both non-target death rates were set to 0.5, all three patterns achieved stability at *D >* 0.7 across all interconversion rates (Figure 7b). Notably, stability was completely independent of interconversion rate in this condition — a striking contrast to the N-suppressing patterns. This is consistent with the drug dose threshold of *D >* 0.7 mapping directly onto the N growth rate of *k*1*_N_* = 0.75; with both non-target populations subject to death rates, the system is sufficiently suppressed that the balance between N growth and drug-induced death alone determines stability, and interconversion between populations no longer influences the outcome.

### 3.4 Analysis of Simulated Hypothetical Treatment Schedules for Validation

Two, three and four-phase treatments were applied over 4-cycle schedules to 200 different initial tumour set-ups. Treatment schedules are described in the methods, with treatment A being two-phase, B and C being three-phase, and D being four-phase (Table 4). Time series plots were made for the average cell populations over each treatment schedule, separated by the initial population makeup.

#### 3.4.1 Two Phase Treatment

The two-phase treatment schedule did not apply an inhibitor. Schedule A applied either P3 or P4, followed by P1. The time series of the average populations for the twophase schedule showed that the initial condition was not a significant factor (Figure 8), agreeing with our predictions that P1 suppresses N cells and P3/P4 suppresses S cells. P1 led to an S-dominant population, which was eradicated by P3/P4 in the following phase, wherein the population became dominated by N cells again. S-suppressing patterns led to N dominance but did not completely eradicate S populations, and S repopulated as N was suppressed in the P1 phase.

**Fig. 8:**
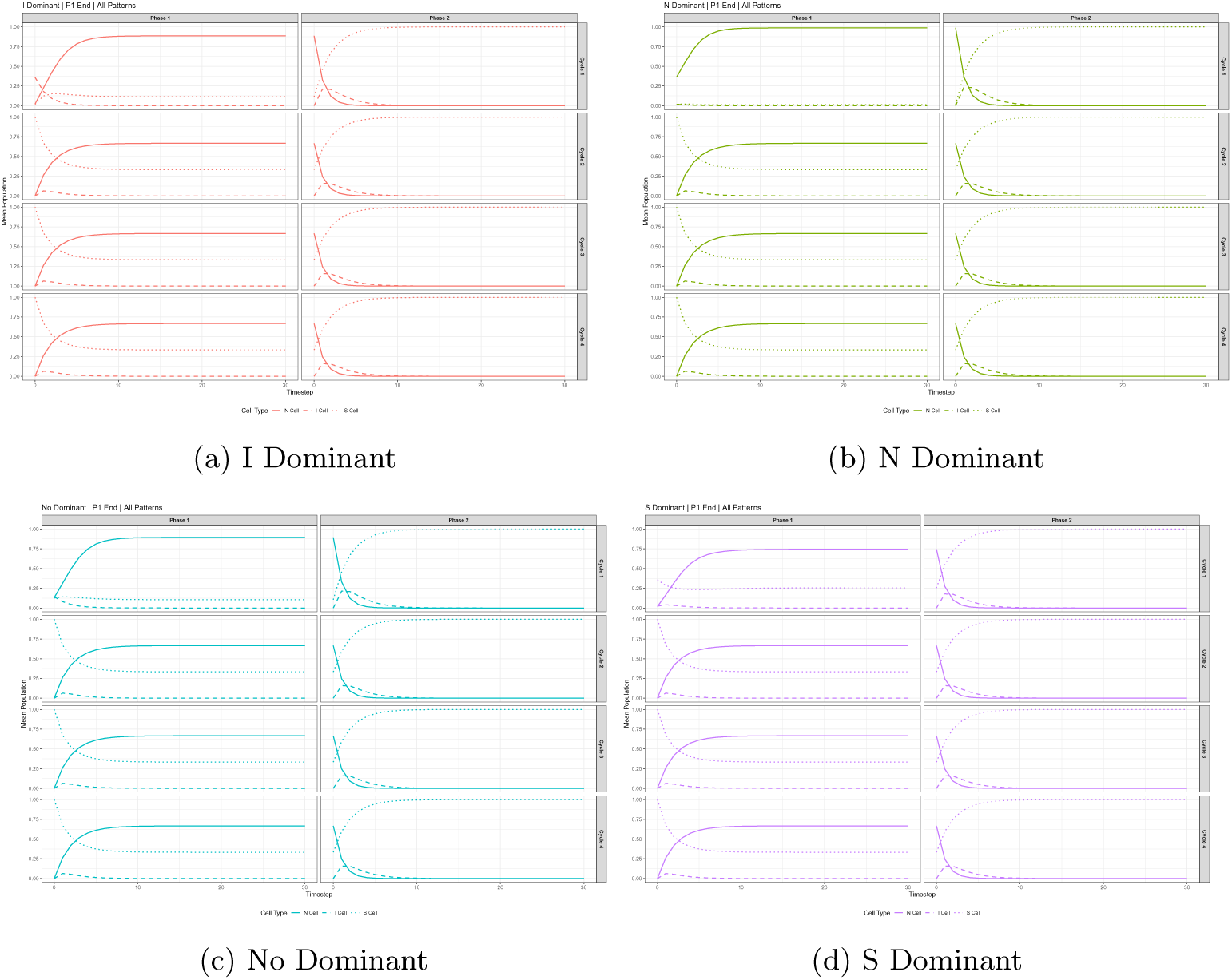
Mean cell population trajectories under the two-phase treatment schedule (Schedule A: one of P3 or P4 in phase one, followed by P1 in phase two, no inhibitor applied), across four treatment cycles. Panels show results faceted by initial tumour composition: (a) I-dominant, (b) N-dominant, (c) no dominant cell type, (d) Sdominant. N (solid), I (dashed), and S (dotted) lines show the mean population over time (days), averaged across all runs for that initial condition.

#### 3.4.2 Three Phase Treatment

Three phase treatment used one inhibitor. Schedule B applied P1, then added an S inhibitor to P1, followed by either P3 or P4. (Figure 9). Schedule C applied P3 or P4, then added an N inhibitor to the pattern, followed by P1 (Figure 10). For treatment B, all initial setups led to a single S cell tumour composition at the end of phase one, which was driven to extinction during phase two from the S inhibitor (Figure 9).The treatment C time series plots show that P3 and P4, on average, are unable to suppress the population into a single cell state during phase one (Figure 10). Thus, although N cells are driven to extinction by the N inhibitor, the S cells remain. This means that, based on the average cell composition over time, full remission is never achieved, although individual cases might. There were no differences in the phase dynamics between cycles 2, 3 and 4. This means that on average, the treatment B was consistent across the different initial cell populations.

**Fig. 9:**
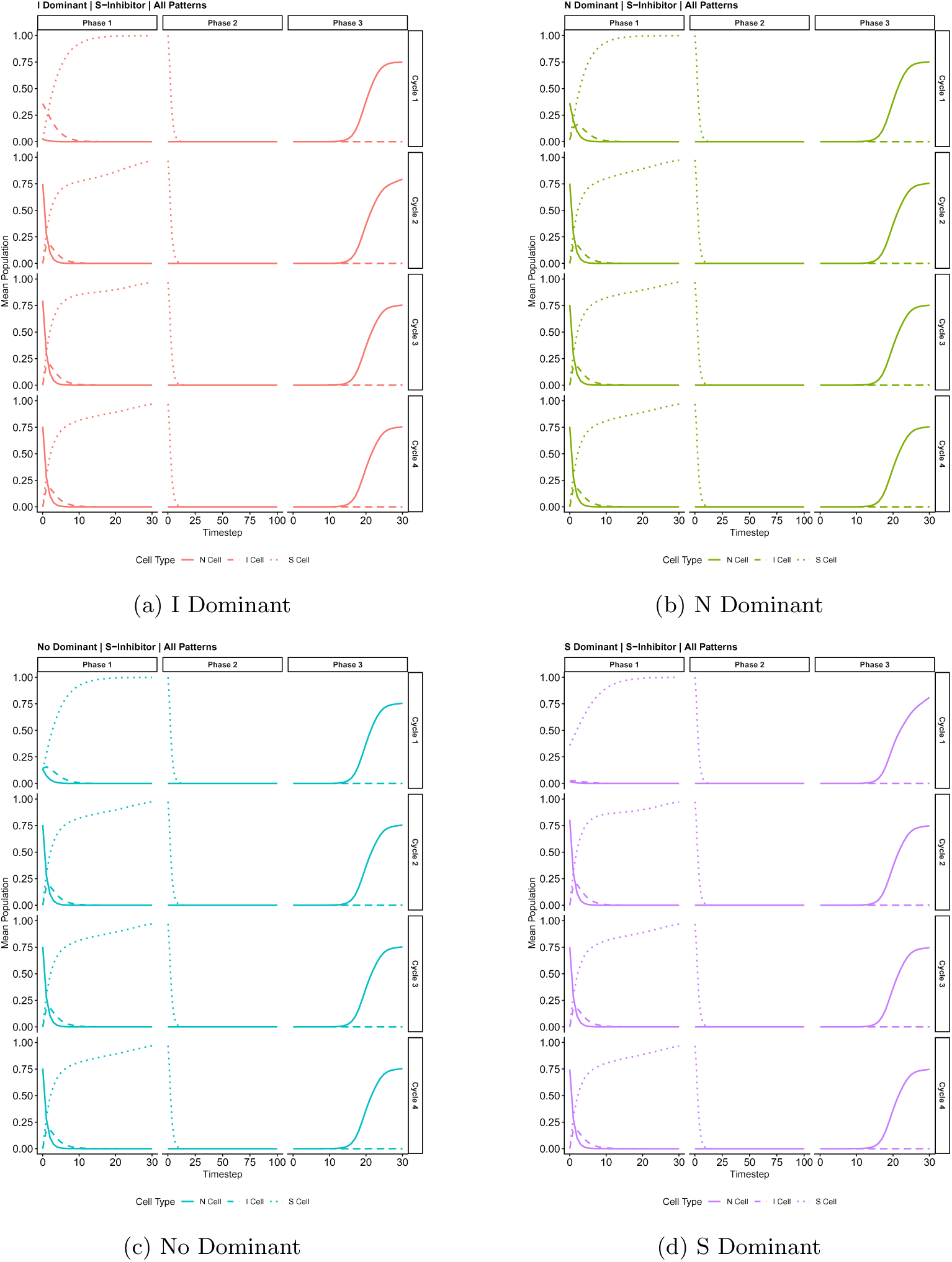
Mean cell population trajectories under the three-phase, S inhibitor treatment schedule (Schedule B: N-suppressing P1, then P1 with S inhibitor, followed by one of P3 or P4 no inhibitor applied), across four treatment cycles. Panels show results faceted by initial tumour composition: (a) I-dominant, (b) N-dominant, (c) no dominant cell type, (d) S-dominant. N (solid), I (dashed), and S (dotted) lines show the mean population over time (days), averaged across all runs for that initial condition.

**Fig. 10:**
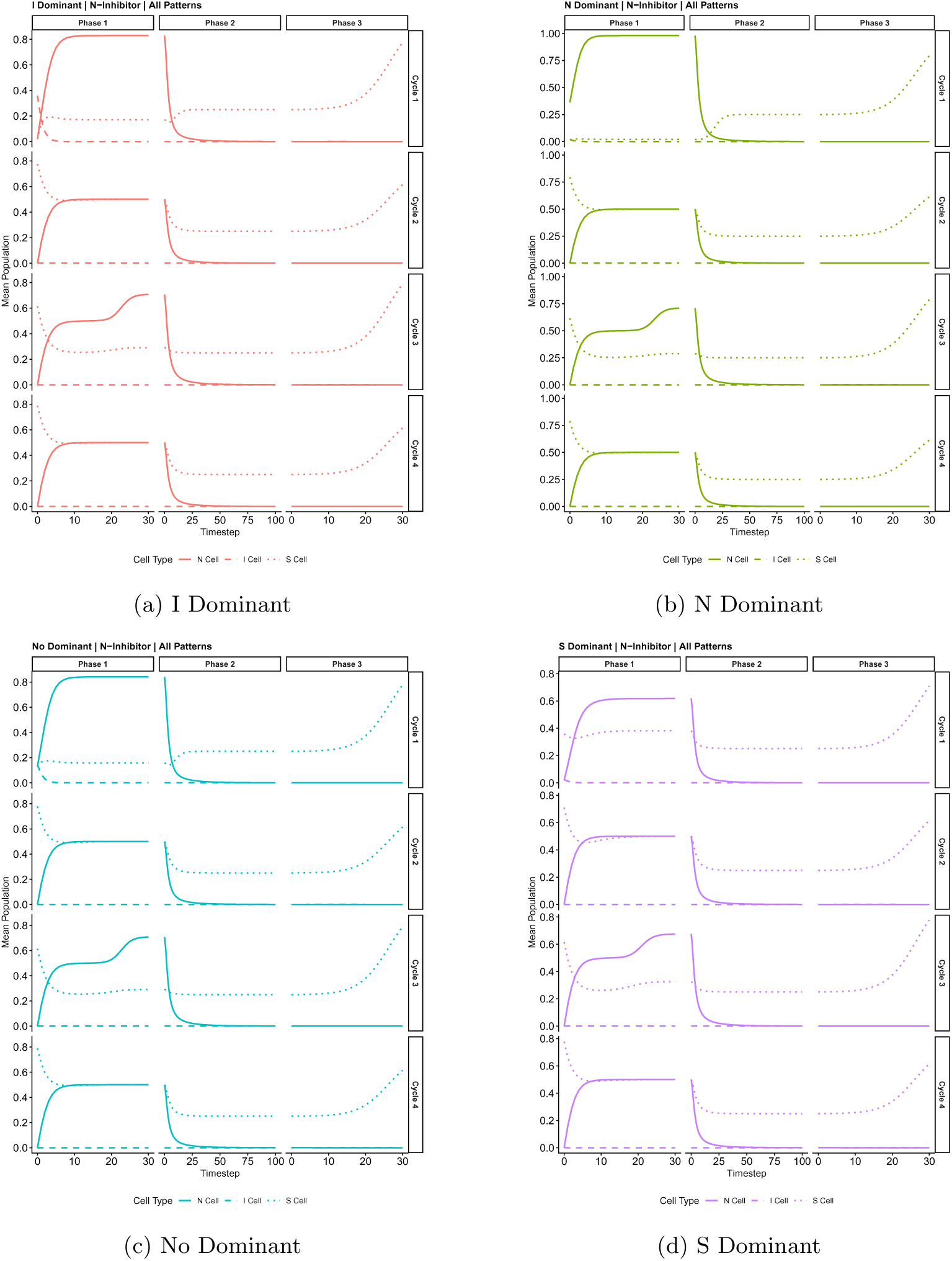
Mean cell population trajectories under the three-phase treatment schedule (Schedule C: S-suppressing P3 or P4, then the same pattern with N inhibitor, followed by P1 no inhibitor applied.), across four treatment cycles. Panels show results faceted by initial tumour composition: (a) I-dominant, (b) N-dominant, (c) no dominant cell type, (d) S-dominant. N (solid), I (dashed), and S (dotted) lines show the mean population over time (days), averaged across all runs for that initial condition.

Comparing the two three-phase treatments, treatment B is the more suppressing order of patterns, as it forces the population into a single cell type, S-dominant first, which with the addition of the S inhibitor in treatment, effectively pushes the total population to extinction and the tumour into remission. Whereas, treatment C, which starts with P3 or P4, does not force the population into a single cell type, and therefore the N inhibitor is unable to force cell extinction. This effect was cause by P4, unlike P3, P4 has an S sink this lead to S cells being present in the final population. This follows the consistent presence of S cells at the end of NTG when P4 was used in the sensitivity analysis and the consistent instability with the absence of *K*2*_S_* = 0.5 in the stability analysis.

#### 3.4.3 Four Phase Treatment

Combining the use of N and S inhibitors in schedule D, Phases 1, 2 and 3 followed the same trajectories as schedule C. Phase 4 was able to push the population to extinction leading to an N-dominant relapse in phase one from cycle 2 onwards. The application on the N inhibitor in phase two actually allowed S cells to repopulate, around 15 timesteps, as the N cell were suppressed (Figure 11), similar to schedule C (Figure 10). Therefore, the addition of the S inhibitor in phase four complements the N inhibitor. Without drug intervention S cells will persist under the suppressing patterns regardless, as seen in treatment A. If only one drug can be applied, schedule B is able to achieve remission with the S inhibitor. Combining both inhibitors in schedule D showed improved outcomes compared to schedule C. Although schedule B and D are equivalent for the application of S in phase two and four respectively, there are fewer known S inhibitors constraining the clinical applications of schedule B as the schedule D can apply a broader repertoire of available N inhibitors. Additionally the combination therapy may offer synergistic effects not captured in this model.

**Fig. 11:**
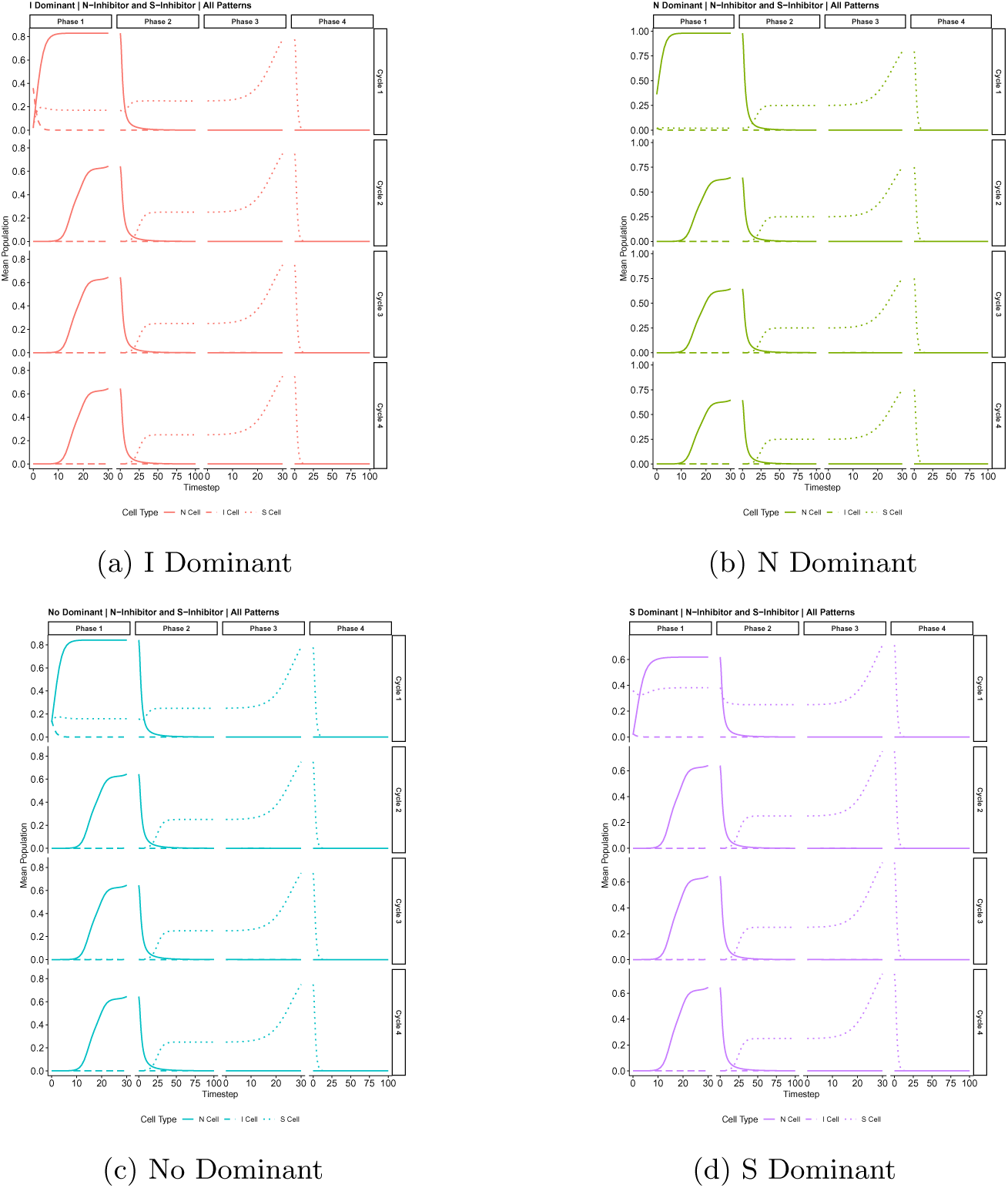
Mean cell population trajectories under the four-phase treatment schedule (Schedule D: S-suppressing P3 or P4, then the same pattern with N inhibitor, followed by P1 no inhibitor applied, then P1 with S inhibitor.), across four treatment cycles. Panels show results faceted by initial tumour composition: (a) I-dominant, (b) Ndominant, (c) no dominant cell type, (d) S-dominant. N (solid), I (dashed), and S (dotted) lines show the mean population over time (days), averaged across all runs for that initial condition.

## 4 Discussion

Intratumoral heterogeneity remains one of the central challenges in treating highrisk neuroblastoma. The mechanisms underlying this heterogeneity are slowly being uncovered, specifically the mechanisms of interconversion between noradrenergic (NOR), mesenchymal (MES) and intermediate (I)-type cells. These cell types can interconvert through core regulatory circuits made up of transcription factors [26, 37]. For NOR, this includes MYCN, PHOX2B, and GATA3, and for MES, PRRX1, NOTCH3, and SOX9 [38, 39]. These cells exhibit resistance to different sets of therapies [9, 14, 40, 41], enabling treatment escape.

Bayesian inference quantified phenotypic adaptations to nutrient stress and found that cell-to-cell interactions drove natural interconversion and adaptation rates [16]. Two cell lines with known initial cell dominance were explored: MES-dominant cells were able to transition to NOR and NOR-dominant cells transitioned to MES; these transitions required a viable mix of both cell types in the initial conditions.

Existing mathematical models have addressed parts of the problem. Using evolutionary game theory as a framework, Bukkuri et al. (2024) model NOR and MES competition under repeated treatment cycles and found that proliferation and resistance better explained behaviour than phenotypic switching [28]. Their model is consistent with P4, where I can switch unidirectionally to N and S, but N and S cannot interconvert. The optimisation of chemotherapy and ALK inhibitor scheduling considering the clonal evolution within neuroblastoma and pharmacokinetics has also been modelled mathematically [29]. Critically, in these models, I-type cells were absent and did not consider phenotypic plasticity. The proposed treatment schedules assume interconversion is negligible. A review of neuroblastoma cell biology identified 34 therapeutically promising interconversion patterns. These patterns are considered to be the most likely naturally occurring, including I-type cells, and to improve clinical management of neuroblastoma [18]. As well as the natural interconversion, compounds such as retinoic acid are known to induce interconversion (Table 9). No model therefore incorporated all three neuroblastoma cell types to optimise treatment as a function of interconversion between cell types; here we aim to fill this gap.

**Table 9:** Summary of known interactions between inhibitors and neuroblastoma cell types. This table compiles experimentally reported relationships between NOR, MES, and I-type cells and the inhibitors that control them. For each cell type, the table lists inhibitors and the mechanistic effect’s direction of inhibition or transition (e.g. NOR→MES, MES→NOR, Itype→MES), along with supporting references.

| Cell Type | Drug / inhibitor | Mechanistic relationship | Reference |
|---|---|---|---|
| NOR | BET inhibitors (JQ1, OTX015) | Inhibit | [40, 41, 46] |
| NOR | ALK inhibitors (lorlatinib, crizotinib) in ALK mutant NOR | Inhibit | [47] |
| NOR | anti GD2 | Inhibit (targets GD2 high NOR cells) Transition: NOR→MES | [48] |
| NOR | Chemotherapy | Inhibit | [9, 14] |
| MES | ALK inhibitor escape | Transition: NOR→MES | [9] |
| MES | MEK inhibitors (in RAS mutant MES) | Inhibit | [49] |
| MES | TRAIL | Inhibit | [9] |
| I-type | Retinoic acid | Transition: I-type→NOR | [3, 50] |
| I-type | BudR | Transition: I-type→MES | [50] |
| I-type | Bmi-1 (Concentration-dependent) | Transition: I-type→NOR | [51] |
| I-type | Bmi-1 (Concentration-dependent) | Transition: I-type→MES | [51] |

A mathematical model was developed; the final product includes NOR, MES and I-type cells to test whether manipulating the interconversion pathways could improve simulated treatment outcomes. The model parameters were grounded in established neuroblastoma biology. NOR-type cells (N) were fastest-proliferating, MES-type (S) cells were slowest-proliferating and I-type cells were assigned an intermediate growth rate. Because interconversion rates between cell types have not been experimentally identified, a range of low, medium, and high values were explored to assess model robustness. Non-target cell death rates were included as known therapeutic targets affect multiple cell types. B7-H3 and L1CAM affect both MES and NOR cells [22], and MEK affects I-type and MES cells [42]. In our model the interconversion of the cells are intrinsic, the dynamics that could induce interconversion are not explored such as drug induced interconversion as see in Table 9 First, we looked at how the 64 possible patterns affect tumour cell populations. We found two N-suppressing patterns (P1 and P2) and three S-suppressing patterns (P3, P4, and P5). P4 is consistent with the competing model. Then we asked which factors of the model affect the shortlisted patterns, including a stability analysis on the origin (remission). Finally, we combined these patterns, an N inhibitor, and an S inhibitor in multi-phase treatment schedules, each of which was deployed in cycles.

The framework presented here, in which a small number of cell states interconvert and differ in treatment sensitivity, such as in breast cancer cell lines [43], small cell lung cancer[44, 45].

### 4.1 In Silico Drug trial

Patterns that suppress either N or S populations in the in silico drug trial were identified. There are five shortlisted patterns. P1 and P2 suppress N cells effectively. P3, P4, and P5 suppress S cells effectively. These shortlisted patterns all had paths for I to interconvert into N or S. On average, a shortlisted pattern reduced the target population to a level below 0.05 in the NTG phase of the in silico trial. Additionally, a shortlisted pattern reduced the I population to a low level in the NTG phase of the in silico trial on average, compared to the other patterns. After the NTG phase, the complementary inhibitor was applied to each shortlisted pattern in the treatment phase. For example, an S inhibitor was applied in conjunction with P1.

P1 (*< N >* |*I >* |*S*|) was observed to be plausible for natural occurrence [18] and is an N-suppressing pattern in which N cells can interconvert to I and S, and I cells can convert to S. No compound has been characterised for the N-to-I transition. However, N-to-S has multiple possible compounds, including RA and the use of an anti GD2 treatment. Likewise, the I-to-S transition can be achieved using BudR and Bmi-1. This pattern is of interest, but more research is needed on compounds that induce the N-to-I pathway. During NTG, N and I are channelled into the S cell type. This traps the faster-growing cell populations in the slower-proliferating cell type. When paired with the S inhibitor, P1 was the most consistent N-suppressing pattern across runs. This is due to the joint mechanisms of cells being converted to a slower-growing type and then targeted by the inhibitor.

P2 (*< N >* |*I >< S*|) was the second N-suppressing pattern and is not considered to be plausible for natural occurrence [18]. For the pattern, S is not acting as a sink because it is able to interconvert to I-type. N cells can interconvert to I and S, and I cells can convert to S. The S to I transition pathway is the least studied, with no known way of inducing this path; it is thought that because I-type cells are undifferentiated [52], it is harder to go from the more common MES type to the intermediate type.

P3 (|*N* | *< I >* |*S >*) is an S-suppressing pattern and is biologically plausible as a naturally occurring process [18]. The I to N transition can be induced by RA, and the I to S pathways can be induced by BudR, though which pathway is activated is concentration-dependent. Retinoic acid (RA) and CDK4/6 have been shown to induce the S-to-N pathway [53].

The second S-suppressing pattern, P4 (|*N* | *< I >* |*S*|), was also consistent with a naturally occurring, biologically plausible process [18]. P4 has both N and S acting as a sink for I-type cells. This produces a different dynamic from P3 or P5: the double sink leads to I cells being suppressed during NTG while N and S accumulate, meaning the initial population conditions can have a large impact. When paired with the N inhibitor, the I-type cells directed to N cell types are eradicated, whereas the I-type cells directed to S are suppressed by their slower proliferation rate but never driven to extinction.

Finally, P5 (|*N* | *< I >< S >*) was an S-suppressing pattern not found to have plausible biological support as a natural occurrence [18]. This pattern sees N as the only sink, the same as P3. However, here S is converting to I, thereby becoming a faster-proliferating cell type. As discussed above, the S to I transition pathway is the least studied and hardest to induce.

Overall, the S-suppressing patterns were on average better at reducing the target cell populations and I-type cells during NTG than the N-suppressing patterns. Additionally, the relative reduction score for the treatment phase was higher for S-suppressing patterns. P2 had the worst relative reduction score for the treatment phase and, alongside P5, was not identified as a plausible naturally occurring transition [18], leading to both patterns being removed for the validation step.

### 4.2 Sensitivity Analysis

The first key finding from the sensitivity analysis was that P1 for N-suppressing and P4 for S-suppressing patterns reduced target cell populations across more runs with medium drug doses compared to the other patterns. This is important because lower doses reduce drug toxicity and side effects [54, 55]. Both patterns have S as a sink, with cells being able to interconvert into S but not away from it. This means for P1, N and I cells are being trapped, by interconversion,into the S cell type and killed under the S inhibitor.

These structural properties are maintained regardless of interconversion rate. For N-suppressing patterns, P1 had the highest relative reduction score across interconversion rates. Compared to S-suppressing patterns, N-suppressing patterns require a medium-to-high interconversion rate, whereas S-suppressing patterns had greater relative reductions at lower interconversion rates. This is because S-suppressing patterns push cells toward the N cell type, which has a faster proliferation rate than S and I, and a lower interconversion rate allows time for N cells to be killed by the corresponding inhibitor. P4 is less sensitive to interconversion rates than P3 and P5, as cells also transition into the slower-proliferating S-type cell.

### 4.3 Stability Analysis

Here, the origin defines remission as extinction of the tumour cell population and being stable defines the maintenance of remission and being robust to perturbation, thereby representing protection against clinical relapse [56, 57]. The origin during NTG was unstable for all five patterns due to the logistic growth rate. It is worth noting that logistic growth prevents small populations from exhibiting dynamics that lead to stochastic extinction, as seen with the Allee effect, where a population’s per-capita growth rate decreases with population density [58]. Alternative common tumour growth models include Mendelsohn, linear, surface, and Gompertz models [59].

For the treatment phase analysis, P1, combined with an S inhibitor, had the most consistently stable origin across interconversion rates and drug concentrations. This is due to the S cell-type sink, leading to cells becoming slower-growing and being eliminated by the S inhibitor. Therefore, P1 plus S inhibitor is the most optimal configuration for the maintenance of remission. As previously discussed, the P1 is a plausible naturally occurring pattern but is also inducible, as RA and anti-GD2 can induce the N-to-S pathway, and the I-to-S transition can be achieved with BudR and Bmi-1. RA is already being tested alongside anti-GD2 and chemotherapy drugs [60–62]. However, when both non-target cell death rates are set to 0.5, the stability of P3, P4, and P5 does not depend on the interconversion rates (Figure 7b).

P3 and P5 dependably showed a bifurcation event at drug concentration 0.7. This shows that the drug needs to be at the N-type cell growth rate so that the cell death rate can outpace the growth rate of the faster-growing N-type cells.

P4’s stability, however, was dependent on the S-type cell death rates. When the Stype cell death rate was set to 0, the lowest eigenvalue of P4 was 0.25, which matched the S cell growth rate; this is due to the S sink seen in the pattern structure and shows how S accumulation destabilised the system. A sufficiently high S cell death rate is needed to allow the drug to reach a stable origin. Non-target cell death rate was set to a binary set of parameters; it is possible that a higher non-target cell death rate could lead to a reduction in drug concentration needed.

### 4.4 Schedule Validation

Because P2 had fewer runs that attained the relative reductions score in the sensitivity analysis and, overall, more unstable origins the tumour cell population. Additionally P2 and P5 are not considered plausible for natural occurrence [18]. Therefore, P2 and P5 were removed from the validity test. The two S-suppressing patterns P3 or P4 were also used with P1, the N-suppressing pattern, in the validity analysis to assess how sequential patterns suppressed cells.

Four schedules deploying P1, P3, P4, an N inhibitor, and an S inhibitor were devised. Each was applied in cycles as seen in clinical treatment [63, 64]; a continuous fixed drug dose was used during a treatment phase.

Without inhibitors, extinction could not be achieved when Schedule A was applied due to logistic growth. However, by switching between P1 and P3/P4, Schedule A resulted in oscillations between N-dominant and S-dominant populations.

While this model uses intrinsic interconversion of the cells, compounds that induce interconversion are already being used alongside treatments. RA induces interconversion and has been tested alongside chemotherapy and a CDk4/6 inhibitor, showing reprogramming is possible to improve targeted therapy [53, 63, 65]. With the addition of one inhibitor, the mean population was more consistent compared to no inhibitor.

Schedule C combined an S-suppressing pattern and an N inhibitor. Schedule C did not show remission in the mean populations. This was because, whilst all P3 runs were able to reach remission, under P4, not all runs reached remission, although those that did, did so faster than under P3. The failure rate was due to the S sink seen in P4. The time to remission was affected by the initial tumour makeup; when initial conditions were I or S dominant, the time to remission was reduced. P4 drives I-types cells toward N and S types, which are suppressed by the N inhibitor and S-type slow proliferation rate. P1 drives the Nand I-type cells toward S and eradicates them with the S inhibitor.

Schedule D applies S-suppressing patterns (P3 or P4), then adds the N inhibitor, then N-suppressing patterns (P1), and finally adds the S inhibitor. Under this schedule, P1 was able to further reduce the time to remission for phase two by suppressing the number of N cells for the N inhibitor to kill. Reducing the amount of time in a drug-induced state is important for lowering toxicity and side effects [54, 55]. The use of two inhibitors has already been applied clinically, such as anti-GD2 with chemotherapy or ALKi and chemotherapy [55, 64].

Overall, inhibitors are necessary; if S inhibitors are available, the P1 pattern followed by S inhibitors is the most optimal for reducing the drug load. Adding the S-suppressing patterns and an N inhibitor can lead to longer remissions under the P1 pattern. Using the S-suppressing patterns and an N inhibitor, the tumour dynamics play a role. P3 achieved remission in all runs, whereas P4 had some scenarios where remission was not achieved. However, when P4 did result in remission, it did so faster than P3.

### 4.5 Research Validation

From these results, the utility of the interconversion patterns is seen. Of the 5 final patterns of interest that reduce S and N populations, three patterns were identified as naturally observed interconversion trajectories by Zeineldin (2022) [18].

Additionally, this model offers an alternative way to identify plausible natural interconversion patterns by looking at cell populations at diagnosis and after treatment. Experimentally, NOR populations were observed at 40 60% at diagnosis and 10% after chemotherapy, whereas MES populations were observed to be at 10% at diagnosis and 10 60% after chemotherapy [23, 28]. By examining the mean cell populations at the end of NTG, and after N inhibitor treatment, we can identify patterns that match these proportions, supporting the biological plausibility of the model output.

Known compounds can be used to induce interconversion pathways [9, 50, 51], and these compounds are already being used alongside therapies to improve patient outcomes [53, 63, 65]. But the viability of the induction of the 64 patterns needs to be explored, and by identifying theoretically optimal patterns, this process can be streamlined. From our 5 patterns of interest, N to S, I to N, and I to S all have known compounds to induce interconversions (Table 9). However, S to N and S to I are not currently inducable.

Model parameters were grounded in established neuroblastoma biology where possible. For the growth rates, N cells were set as the fastest proliferating to mimic NOR [16, 17]. As experimentally measured values are unavailable for interconversion rates and death rates, clinically relevant variability was used to give a reasonable range. The initial tumour cell makeup is also known to vary depending on cell line [4, 11, 12, 22]. Therefore, for the validation of the model a different cell dominance initial condition was used.

Cell dynamics, including growth rate and non-target cell death rate, affect the schedule efficacy, which means the initial tumour cell makeup is a key piece of information. Imaging can give initial tumour makeup to optimise treatment. Molecular profiling and histology can be used for therapeutic stratification [26, 36, 37, 66, 67]. A combination of image and single-cell sequencing data can be used to identify what cells are dominant in the tumour, and with an AI tool can identify the optimal treatments [67–71].

### 4.6 Limitations

This model uses logistic growth for the three cell types, which meant that remission without drug is not possible. An alternative would be to simulate the Allee effect, where small tumour cell populations undergo stochastic extinction [58]. Other growth models are being explored [59]. The model has no spatial component, meaning neighbourhood effects and cell-to-cell interactions cannot be investigated with this model; it is a major limitation because these interactions promote interconversion [16].

The model ignores genetic mutations. DNA methylation, histone modifications, and chromatin components can lead to resistance in cancers [72, 73]; these epigenetic changes will persist in the cell even if it interconverts into a different cell type [74–76].

The schedules we tested were limited to a continuous fixed-dose delivery of one drug per phase. Real schedules often use different intervals depending on the half-life of the drug being used, and multiple drugs may be induced at the same time [77–80]. Additionally, optimisation was not systematic; as schedules become more complex, a systematic approach would allow for optimal schedules to be discovered. The combination of tumour imaging and single-cell sequencing will allow for realistic initial cell composition to optimise model calibration.

Future work should incorporate drug-dependent interconversion terms, enabling the model to better fit clinical cell data and combination treatment as described in Table 9. Additionally, applying different treatment schedules, with two inhibitors applied simultaneously, and different dosing schedules, will allow for a more realistic simulation of clinical treatment. Finally, the model should be extended to include resistance mutation to capture the dynamics of treatment response.

## 5 Conclusion

We developed a three-population model of noradrenergic (NOR), mesenchymal (MES) and intermediate (I-type) neuroblastoma cells to ask how the pattern of interconversion between these cell types affects phenotype-specific inhibitors. Of the 64 possible patterns, three patterns suppressed their target population during natural tumour growth and have been reported as biologically plausible: one for N cells and two for S cells. N-suppressing patterns worked best at medium-to-high interconversion rates, and S-suppressing patterns at lower rates. During treatment, an N-suppressing pattern combined with an S inhibitor gave the most consistently stable tumour-free state. Schedules that alternate Nand S-suppressing patterns with the matching inhibitor suppressed the tumour more reliably than single-pattern treatment.

These results are from a deterministic model with uncalibrated parameters, logistic growth, no spatial structure and no resistance mutations, so they identify candidate strategies rather than treatment recommendations. Future work should add drugdependent interconversion terms, calibrate the model to single-cell and imaging data, and optimise treatment schedules systematically. Because the framework uses a number of interconverting cell states, it may also apply to other cancers with documented plasticity, such as breast cancer and small cell lung cancer.

## Acknowledgement

KYW conceived the project. KYW and MP secured funding for the project. FC, KYW, and MP designed the study presented in this manuscript. FC implemented the design by constructing the mathematical model, parameterising it, conducting computer simulations, and carrying out a stability analysis. FC analysed and interpreted the results. FC created the visualisations presented in the manuscript. FC wrote the manuscript. FC, KYW, and MP reviewed and edited the manuscript. KYW and MP provided scientific leadership by supervising every step of the study. All three authors have read and approved the manuscript. FC is grateful for a PhD studentship awarded by the Centre of Excellence for Data Science, Artificial Intelligence, and Modelling, the University of Hull. FC, KYW, and MP are grateful for Dr Sabine Taschner-Mandl’s scientific advice.

## Statements and Declarations

### Data and code availability

All code and data are available at https://github.com/frcovell/NB Phenotypic Plasticity and archived on Zenodo (https://doi.org/10.5281/zenodo.23744299).

